# Microglial P2Y12 mediates chronic stress-induced synapse loss in the prefrontal cortex and associated behavioral consequences in male mice

**DOI:** 10.1101/2022.03.16.484487

**Authors:** J.L. Bollinger, D.T. Dadosky, J.K. Flurer, I.L. Rainer, S.C. Woodburn, E.S. Wohleb

**Affiliations:** Department of Pharmacology & Systems Physiology, University of Cincinnati College of Medicine, Cincinnati, OH

## Abstract

Recent studies demonstrate that chronic unpredictable stress (CUS) drives microglia-mediated neuronal remodeling, contributing to synapse loss in the prefrontal cortex (PFC) and cognitive-behavioral dysfunction. Nonetheless, it remains unclear what mechanisms guide microglia-neuron interactions in stress. Evidence indicates that neuronal activity-dependent purinergic signaling directs microglial processes and microglia-synapse interaction via P2Y12, a purinergic receptor exclusively expressed by microglia in the brain. Stress exposure alters excitatory neurotransmission in the PFC, thus we aimed to determine if P2Y12 signaling promotes functional changes in microglia in the context of chronic stress. Using an activating DREADD, our initial studies showed that PFC microglia adopt a CUS-associated phenotype after repeated pyramidal neuronal activation. To further investigate the role of purinergic signaling, we used genetic (*P2ry12-/-*) or pharmacological (clopidogrel, ticagrelor) approaches to block P2Y12 in the context of CUS. Various behavioral, physiological, and cytometric endpoints were analyzed. Both P2Y12-deletion and treatment with clopidogrel prevented increases in forced swim test immobility and attenuated deficits in temporal object recognition following CUS. Flow cytometry of PFC microglia revealed that both *P2ry12-/-* mice and those treated with clopidogrel have significantly different phenotypes (independent of CUS); with diminished P2Y12 expression and altered surface levels of CX3CR1, CSF1R, and CD11b. Immunohistology in Thy1-GFP(M) mice demonstrated that pharmacological blockade of P2Y12 prevented stress-induced increases in the proportion of microglia with GFP+ neuronal inclusions and limited dendritic spine loss in the PFC. Together, these findings indicate that microglial P2Y12 is a critical mediator of stress-induced neuronal remodeling in the PFC and subsequent behavioral deficits.

## Introduction

Alterations in synaptic plasticity are thought to underlie numerous stress-linked psychiatric disorders, including major depressive disorder (MDD). For instance, patients suffering from MDD have fewer synapses in the prefrontal cortex (PFC) and this correlates with MDD severity and working memory dysfunction [1]. Clinical and preclinical findings suggest that microglia, the brain-resident macrophages, may contribute to synaptic alterations in the PFC and MDD-associated symptoms. In fact, chronic stress has been shown to induce microglia-mediated remodeling of synapses in the medial PFC (mPFC) of mice and subsequent deficits in PFC-mediated behaviors, including working memory function and cognitive flexibility [2, 3]. Despite these findings, relatively little is known regarding the pathways that drive microglia-neuron interactions in stress and microglial interventions in stress-linked disorders remain elusive.

Studies indicate that initial stress exposure increases pyramidal neuron activity and disturbs neuronal homeostasis in the PFC [4, 5]. Microglia respond dynamically to changes in neuronal activity and the local release of purines, including adenosine-diphosphate and triphosphate (ADP and ATP) [6, 7]. Purine release by neurons attracts microglial processes via P2Y12, a purinergic receptor exclusively expressed by microglia in the brain [8–10]. These activity-dependent interactions have functional consequences as mice with genetic deletion of P2Y12 (*P2ry12-/-*) show impaired plasticity in the visual cortex following monocular deprivation during neurodevelopment; and this was attributed to reduced uptake of synaptic material by microglia [8]. Thus, it is plausible that P2Y12 plays an important role in guiding stress-induced changes in microglia-mediated synaptic remodeling in the mPFC and associated behavioral consequences. To investigate this, we first examined microglial phenotype in response to protracted neuronal activation in the mPFC using a DREADD-mediated approach. We next disrupted P2Y12 signaling during chronic stress using either genetic or pharmacological tactics, with analyses addressing microglia-neuron interactions in the mPFC, synapse density in this region, stress physiology, and cognitive-behavioral function.

## Materials and Methods

### Animals

Transgenic *P2ry12-/-*, Thy1-GFP(M), and floxed-hM3Dq mice were obtained from in-house breeders backcrossed to C57BL/6J mice (Jackson Laboratories; C57BL/6J; #000664). Transgenic P2Y12 breeders were generously provided by Dr. Ania Majewska (University of Rochester Medical Center, Rochester, NY), both floxed-hM3Dq (RC::L-hM3Dq, #026943) and Thy1-GFP(M) (Tg(Thy1-EGFP)MJrs/J, #007788) breeders were acquired from Jackson Laboratories. Thy1-GFP(M)/P2Y12 transgenic mice were generated by crossing lines. Complementary clopidogrel and ticagrelor experiments were performed using Thy1-GFP(M) and wild-type C57BL/6J mice. All mice (6-8 weeks old) were group-housed (3-4/cage) in 11.5×7.5×6 inch polypropylene cages under a 12-hour light-dark cycle with ad libitum access to food and water. Previous studies indicate microglia-mediated neuronal remodeling in the PFC in stressed male-but not female-mice [3, 11]. Therefore, all experiments were conducted in male mice, with only initial behavioral- and physiological-analyses examined in females lacking P2Y12. Animal procedures were approved by the University of Cincinnati Institutional Animal Care and Use Committees (IACUC) and were in accordance with guidelines established by the National Institutes of Health.

### Chronic unpredictable stress (CUS)

CUS was performed as previously described [3, 12]. In brief, mice were exposed to two random intermittent stressors per day for 14 days, including: cage rotation, isolation, restraint, radio noise, food or water deprivation, light on overnight, light off during day, rat odor, stroboscope overnight, crowding, wet bedding, no bedding, or tilted cage. This paradigm increases circulating corticosterone, induces adrenal hypertrophy, and reduces animal weight gain [12].

### Drug administration

Clozapine N-oxide (CNO, National Institutes of Health, Bethesda, Maryland) was diluted to 0.6 mg/mL in dimethyl sulfoxide (DMSO):0.9% sodium chloride (1:20 ratio). Clopidogrel sulfate (LKT Laboratories, St Paul, MN; C4658) was diluted to 5 mg/ml in 0.9% sodium chloride. Ticagrelor (LKT Laboratories; T3200) was diluted to 6 mg/ml in DMSO:0.9% sodium chloride (2:1 ratio). Mice received either CNO (1 mg/kg), clopidogrel (50 mg/kg), or ticagrelor (10 mg/kg) via daily intraperitoneal (i.p.) injection [8]. Control mice received equal volume injections of 0.9% sodium chloride or DMSO:0.9% sodium chloride. Injections were performed in the morning prior to daily stressors or after behavioral testing in specified cohorts.

### Stereotactic surgery & chemogenetic manipulation

For these experiments, floxed-hM3Dq mice were deeply anesthetized with ketamine:xylazine (90 mg/kg:20 mg/kg) followed by stereotactic surgery. AAV5-CaMKII-mCherry-Cre (UNC Vector Core, Chapel Hill, NC) was bilaterally infused (0.5 μl; 0.1 μl/min) into the mPFC (AP: +2.0mm, ML: ±0.2mm, DV: -2.8mm). Syringes were left in place for 5 min after infusion to limit viral spreading in the needle tract. Mice received daily injection of meloxicam (5 mg/kg, i.p.) over 3 days for pain relief. Mice were handled intermittently over three weeks to allow for surgical recovery and viral infection/expression. Following recovery mice were administered vehicle or CNO for 14 days.

### Behavior and cognitive tests

Forced swim test (FST) and temporal order recognition testing (TOR) were conducted as previously described [3, 12, 13]. All behavioral tests were performed in a normally lit room (white light), between 0800 and 1200 hours. Mice were allowed to habituate to this room for 30 min prior to testing. For the FST, mice were placed in a 2-liter beaker of water (24°+/- 1°C) for 8 min and time spent immobile (2-6 min phase) was measured. In the first trial phase of the TOR, mice were placed in a plastic arena with two plastic Lego™ trees secured to the bottom of the arena. For each phase of the TOR, mice were given 5 min to explore the arena and objects. After a 30 min latency, mice were placed in the same arena with Lego™ blocks (second trial phase). An hour after the second trial phase, mice were placed in the arena with one block and one tree, counterbalanced to prevent bias (test phase). The time spent exploring each of these objects was measured and a discrimination index was calculated (difference between time spent exploring the tree or the block divided by the total time spent exploring the tree and the block). Behavioral testing occurred on either day 13 (FST) or 14 (FST, TOR) of CUS or one day post-CUS (day 15, TOR). FST and TOR were counterbalanced across days 13 and 14 to reduce any effect of prior behavioral testing on performance.

### Stressor validation and physiological stress responsivity

To verify stressfulness of the manipulation, body weight was measured prior to day 1- and on day 14-of CUS, with percent weight gain calculated for each animal for all experimental groups. The following analyses were conducted in *P2ry12-/-* studies. At 2 h post-behavioral testing, a subset of CUS-exposed mice were restrained for 1 h to assess acute stress effects on plasma corticosterone. Immediately following restraint, blood from both CUS animals and unstressed, control animals was collected and stored on ice. Blood was centrifuged at 2000g for 15 min, with plasma collected and stored at -20°C. Circulating corticosterone was then measured via radioimmunoassay (MP Biomedicals, Irvine, CA). In addition to blood collection, animals were euthanized and both adrenal glands and spleen were collected and weighed. Organ-weight-to-body-weight ratios were calculated and compared across groups.

### Percoll gradient enrichment of microglia and flow cytometry

Dissected frontal cortex (cortex rostral to the optic chiasm) was passed through a 70 μm cell strainer. Homogenates were centrifuged at 800g for 8 min. Supernatant was then removed and cells were enriched using one of two Percoll-based approaches (GE Healthcare, Uppsalla, Sweden, #17089102). For DREADD experiments, cell pellets were re-suspended in 30% isotonic Percoll and layered with 0% isotonic Percoll (phosphate buffered saline, PBS). For all other experiments, cell pellets were re-suspended in 70% isotonic Percoll with a discontinuous density gradient layered as follows: 30% and 0% isotonic Percoll. Cell suspensions were centrifuged for 20 min at 2000g. Cells were either pelleted (DREADD experiments) or enriched microglia were collected from the interphase between the 70% and 30% Percoll layers (all other experiments). Fc receptors were then blocked with an anti-CD16/CD32 antibody (BioLegend, San Diego, CA, U.S.A., #553141). Cells were washed and then incubated with conjugated antibodies (FITC-CX3CR1, #149020 or AlexaFluor488-CD115, #135512; PE-P2Y12, #848004 or PE-CD68, #137014; Biolegend; PerCp-Cy5-CD11b; #550993; PE(594)-CD45, #562420; BD Biosciences, Franklin Lakes, NJ) for 45 min at 4°C. Cells were washed and re-suspended in fluorescence activated cell sorting (FACS) buffer. For DREADD experiments, an average of 26,982±2805 microglia were collected per sample based on CD11b/P2Y12 expression using a BioRad S3e four-color cytometer/cell sorter (Hercules, CA, U.S.A.). For all other experiments, an average of 25,434±971 microglia were examined per sample based on CD11b/CD45^lo^ expression. Data were analyzed using FlowJo software (Ashland, OR, U.S.A.).

### RNA isolation and quantitative real-time PCR

For DREADD experiments, RNA was extracted from microglia using a Single Cell RNA purification kit (Norgen Biotek Corp., Thorold, Canada, #51800). Samples were reverse transcribed, and quantitative real-time PCR was conducted as previously described [12]. Primer sequences are listed in Table.S1.

### Immunohistology

Mice were transcardially perfused with PBS followed by 4% paraformaldehyde (PFA) approximately 4 h after the final stressor or 2 h after behavioral testing. Brains were post-fixed in 4% PFA for 24 h and incubated in 30% sucrose until cryoprotected. Brains were then rapidly frozen and sectioned on a Leica CM2050 S cryostat (40 μm). Free-floating sections containing the mPFC were selected for analysis. Sections were washed, blocked in 1% bovine serum albumin (BSA; Fisher Scientific, Waltham, MA; #BP9703) with 5% normal donkey serum (NDS; EMD Milipore, Billerica, MA; #S30-100ML) for 1 h at room temperature, washed, and then incubated with primary antibody: rabbit anti-IBA1 (1:1000, Wako; 019-19741), rabbit anti-P2Y12 (1:500, Anaspec; AS-55043A), rat anti-CD68 (1:500, Bio-Rad; MCA1957), or rabbit anti-FOSB (1:500, Abcam; ab184938), overnight at 4ºC. Sections were then washed and incubated with conjugated secondary antibody overnight at 4ºC (1:1000, Invitrogen; Alexa Fluor 488; Alexa Fluor 546; Alexa Fluor 647). Sections were subsequently washed, mounted on gel-coated slides, and coverslipped with Fluoromount-G (ThermoFisher Scientific; 00-4958-02).

### Quantitative immunofluorescence

Confocal images were obtained on a Nikon C2+ microscope interfaced with a Nikon C2si+ camera (Tokyo, Japan). Confocal images were captured from adjacent brain sections in both hemispheres of the mPFC (3-4 sections/sample). FosB immunolabeled tissue sections were imaged using a 10× objective (NA: 0.95, z-stack: 0.8 μm, image size: 1024×1024). To quantify FosB+ cells, bright immunolabled cell bodies were thresholded and then counted using the ImageJ (NIH) Analyze Particles function (circularity: 0.2-1, size: 30-300). FosB+ cell density was calculated as cells per mm^2^. For analysis of IBA1+ material and P2Y12 intensity, tissue sections were imaged using a 20× objective (NA: 0.95, z-stack: 0.8 μm, image size: 1024×1024). Individual IBA1+ and P2Y12+ cell counts were obtained by hand and cell density was calculated (cells per mm^2^). For microglial area, images were thresholded, total area was recorded (μm^2^), and area per microglial cell was calculated as total area/total number of IBA1+ cells per image (μm^2^/cell). Microglial clustering was assessed using the nearest neighbor distance plugin. For analysis of microglial P2Y12 expression, images were thesholded, fluorescence intensity was measured (integrated density, A.U.), and P2Y12 expression per microglial cell was calculated as fluorescence intensity/total number of IBA1+ cells per image (A.U., relative to control).

In Thy1-GFP(M) mice, layer I of the mPFC was identified in multiple adjacent sections and apical dendritic segments were imaged using either a 40x (NA: 0.95) or 60x objective (NA: 1.27, zoom: 2x, z-stack: 0.2 μm, image size: 1024×1024). Given the expression profile of GFP in this transgenic line, apical segments in layer I most likely originate from pyramidal cells in layer V and VI of the mPFC [14]. Dendritic segments were traced and measured in NeuronStudio with spines counted by hand (6-8 segments totaling 146.5±2.9 μm/sample). To quantify GFP+ inclusions (total number and volume), confocal images were obtained in multiple adjacent sections and microglia with complete morphological profiles were identified (17.3±0.5 cells/sample). Confocal images were examined in 3-dimensional space and as orthogonal z-stacks using Nikon Elements Image Analysis software. Confocal imaging with these settings provides sufficient resolution to detect synaptic inclusions (average inclusion diameter: 0.25 μm; Weinhard et al., 2018). In addition to GFP+ inclusions, the total volume of microglial CD68 was measured in these images, with data expressed as CD68 volume/total number of IBA1+ cells (μm^3^/cell). Of 629 analyzed GFP+ inclusions, 619 (98.4%) colocalized with immunolabeled CD68.

To examine autofluorescent lipid inclusions (e.g., lipofuscin), microglia were visualized in *P2ry12-/-* mice lacking that Thy1-GFP(M) transgene with images collected as described above. Laser settings were maintained across studies. Autofluorescent inclusions were imaged at 488 nm excitation with an emission spectra of 510 – 574 nm. Images were thresholded to capture the entire volume of each inclusion and fluorescence intensity was measured using Nikon Elements Image Analysis software. The mean fluorescence intensity was averaged across 15.1±1.0 inclusions per animal.

### Statistical analysis

Data were analyzed using GraphPad Prism 8.1.2 (La Jolla, California). Student’s t-tests were used to analyze DREADD experiments (vehicle vs. CNO). For subsequent experiments, significant main effects and interactions were determined using two-way ANOVA (stress×genotype, stress×drug treatment). Differences between group means were evaluated using Sidak’s multiple comparisons test. For pharmacological experiments comparing clopidogrel and ticagrelor, animals treated with either 0.9% saline or DMSO:saline did not differ across any metrics and were thus collapsed into one vehicle-treated group for analysis. Pearson correlation coefficients were computed and analyzed for select variables. The number of animals examined in each analysis is noted in Table.S2, with complete omnibus statistics reported in Table.S3.

## Results

### Repeated neuronal activation shifts microglial phenotype in the frontal cortex and disrupts working memory function

Bilateral infusion of AAV5-CamkIIa-mCherry-Cre enabled selective expression of the hM3Dq DREADD in PFC pyramidal neurons. After 3 weeks of recovery, mice were injected daily with vehicle or CNO and then behavioral, cytometric, and molecular outcomes were assessed (Fig.1A-M). Cognition and behavior were assessed via FST immobility, object exploration, and performance in the TOR task. Treatment with CNO did not alter weight gain (Fig.S1). Likewise, vehicle- and CNO-treated mice spent a similar amount of time immobile in the FST and exploring objects in the TOR (Fig.1C-D). However, chronic administration of CNO disrupted object discrimination in the TOR (t_(18)_=2.78, *p*=0.01, Fig.1E). This suggests that protracted neuronal activation in the PFC is sufficient to disrupt prefrontal function and influence cognitive performance.

**Figure 1.**
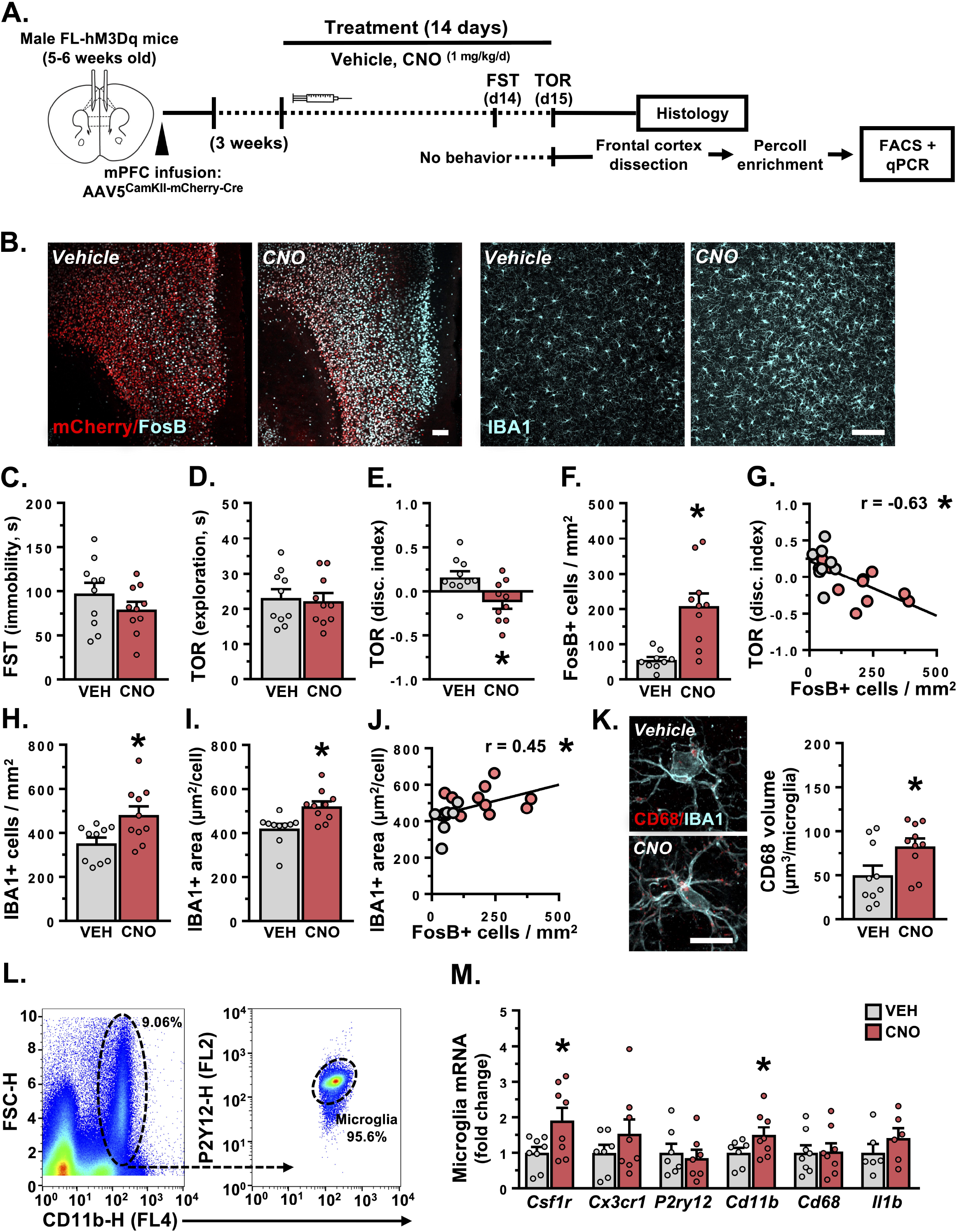
Prolonged prefrontal activation disrupts working memory and shifts microglial phenotype. **A**. Male hM3Dq-floxed mice received bilateral infusions of AAV5-CaMKIIa-mCherry-Cre into the mPFC. Post-recovery, animals received daily injections of either vehicle or CNO for 14 days. One cohort was subjected to behavioral testing, after which mice were perfused and brains were processed for confocal imaging (*n =* 9-10/group). **B**. Left: Representative images of viral mCherry expression (red) and neuronal activation (FOSB, cyan) in the mPFC in vehicle and CNO treated animals. Right: Images of microglia (IBA1, cyan) in the mPFC in vehicle and CNO treated animals. White scale bar represents 100 μm. **C**. Average time spent immobile in the forced swim test (FST). **D**. Average time spent exploring objects in the temporal object recognition task (TOR). **E**. Discrimination index in the TOR. **F**. Average number of FosB+ cells/mm^2^ in the mPFC. **G**. Association between discrimination index in the TOR and the average number of FosB+ cells/mm^2^ in the mPFC. **H**. Average number of IBA1+ cells/mm^2^ in the mPFC. **I**. Average IBA1+ area per microglial cell in the mPFC. **J**. Association between IBA1+ area and the average number of FosB+ cells/mm^2^ in the mPFC. **K**. Left: Confocal images of microglial (cyan) CD68 (red) in the mPFC in vehicle and CNO treated animals. Right: Average volume of CD68 per microglial cell. White scale bar represents 10 μm. **L**. In a separate cohort of mice, brains were extracted, frontal cortex was dissected out, and microglia were isolated via FACS. Microglial gene expression was then characterized (*n =* 6-8/group). Representative dot plots of FSC-H/CD11b-H and P2Y12-H/CD11b-H are shown. **M**. Normalized gene expression of Csf1r, Cx3cr1, P2ry12, Cd11b, Cd68, and Il1b in frontal cortex microglia. Bars represent mean ± S.E.M. ^*^ *p*<0.05 compared to vehicle treated group.

Immunohistology showed that mice treated with CNO had an increased number of FosB+ cells/mm^2^ in the mPFC (t_(17)_=4.01, *p*=0.0009), with heightened FosB induction being correlated with greater dysfunction in the TOR (*r*=-0.625, *p*=0.004; Fig.1F-G). Administration of CNO also increased microglial density (IBA1+ cells/mm^2^; t_(18)_=2.66, *p*=0.02), microglial clustering as assessed using nearest neighbor analysis, and microglial morphological area (IBA1+ area/cell; t_(18)_=3.27, *p*=0.004) in the mPFC (Fig.1H-I, Fig.S1). Heightened levels of FosB were further associated with greater increases in microglial area (*r*=0.425, *p*=0.05; Fig.1J). In addition to shifts in density and morphology, CNO increased the volume of CD68+ lysosomes per microglia in the mPFC (t_(18)_=2.66, *p*=0.02; Fig.1K). Increases in microglial clustering and CD68+ lysosome volume were associated with greater deficits in the TOR (Fig.S1). Together these data indicate an activity-dependent relationship between neuronal function and microglia phenotype in the PFC.

In a separate cohort, microglia were isolated from the frontal cortex using FACS and microglial gene expression was analyzed (Fig.1L). Repeated administration of CNO increased levels of *Csf1r* (t_(14)_=2.38, *p*=0.03) and *Cd11b* (t_(14)_=2.16, *p*=0.05) expression in frontal cortex microglia, yet had little effect on levels of *Cx3cr1, P2ry12, Cd68*, or *Il1b* (Fig.1M). Similar results were observed using AAV5-CaMKIIa-hM3d in the PFC as microglia showed increased *Csf1r* expression after repeated CNO administration (Fig.S2). However, there appears to be a gene-dose response as this manipulation did not cause behavioral or cognitive changes (Fig.S2). This suggests that protracted neuronal activation modulates specific microglial pathways including those relevant to microglia-mediated synaptic remodeling.

### Genetic loss of P2Y12 attenuates stress-induced behavioral consequences and induces significant alterations in frontal cortex microglia

To determine the role of P2Y12 in stress-induced microglia responses, wild-type and *P2ry12-/-* mice were exposed to 14 days of CUS and then physiological, behavioral, and molecular outcomes were assessed (Fig.2A). Consistent with prior studies, CUS increased immobility in the FST (F_(1,40)_=7.78, *p*=0.008), with post-hoc comparisons indicating greater immobility in wild-type (*p*=0.04) but not *P2ry12-/-* mice (Fig.2B). Both unstressed and CUS exposed mice spent a similar amount of time exploring objects in the TOR, regardless of genotype (Fig.2C). However, CUS reduced object discrimination in the TOR (F_(1,40)_=23.83, *p*<0.0001), and these deficits were significant in wild-type mice only (*p*<0.0001; Fig.2D). Post-behavioral testing, we examined whether P2Y12 ablation disrupts physiological stress responsivity. Stressed animals were subjected to a 1 h acute restraint challenge followed by analysis of plasma corticosterone, body weight, and organ weight. Both wild-type and *P2ry12-/-* mice had elevated plasma corticosterone levels after acute restraint (F_(1,38)_=57.34, *p*<0.0001; Fig.2E). Post-hoc findings indicate that stress-associated levels of plasma corticosterone were greater in *P2ry12-/-* mice (*p*=0.04). Additional analyses show that CUS reduced weight gain and induced adrenal hypertrophy in mice regardless of genotype (Fig.S3). Parallel analyses in female mice found no effects of stress or genotype on behavior, despite CUS-induced alterations in weight gain and stress-sensitive organ weight (Fig.S4). Together, these findings indicate an attenuated behavioral response to stress –despite a pronounced physiological response– in male *P2ry12-/-* mice.

**Figure 2.**
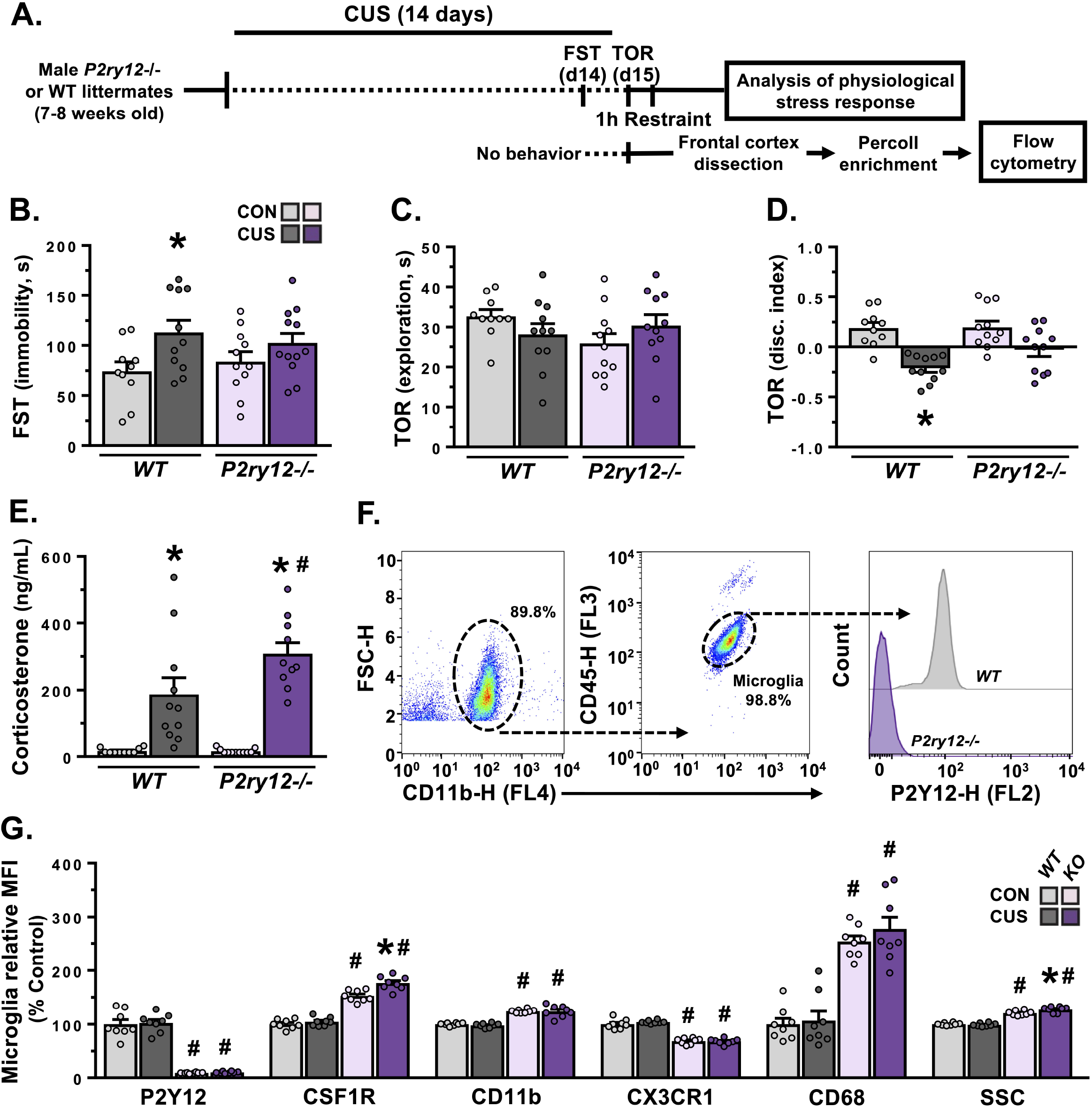
Constitutive ablation of P2Y12 attenuates stress effects on coping behavior and working memory, and induces significant alterations in frontal cortex microglia. **A**. Male wild-type or *P2ry12*-/- mice were exposed to 14 days of chronic unpredictable stress (CUS) or were handled as controls (CON). One cohort was subjected to behavioral testing, an acute stress challenge, and analysis of stress-sensitive organ weight (*n =* 10-12/group). **B**. Average time spent immobile in the forced swim test (FST). **C**. Average time spent exploring objects in the temporal object recognition task (TOR). **D**. Discrimination index in the TOR. **E**. Levels of plasma corticosterone in mice subjected to an acute restraint stress challenge. **F**. In a separate cohort of mice, brains were extracted, frontal cortex was dissected out, and microglia were isolated and characterized using flow cytometry (*n =* 8/group). Representative dot plots of FSC-H/CD11b-H and CD45-H/CD11b-H are shown. Histogram depicts levels of P2Y12 fluorescence in frontal cortex microglia in representative WT and *P2y12-/-* mice. **G**. Normalized mean fluorescence intensity of P2Y12, CSF1R, CD11b, CX3CR1, and CD68 in frontal cortex microglia. Normalized mean side scatter profile is shown. Bars represent mean ± S.E.M. ^*^ *p*<0.05 compared to same-genotype unstressed animal. ^#^ *p<*0.05 compared to unstressed wild-type animal.

In subsequent experiments, surface markers were characterized on frontal cortex microglia using flow cytometry (Fig.2F-G). Analyses of cell surface expression confirmed the absence of P2Y12 in *P2ry12-/-* mice (F_(1,28)_=295.0, *p*<0.0001; Fig.2F&G). Further panels revealed that frontal cortex microglia from *P2ry12-/-* mice had significantly different phenotypes. In particular, microglia had increased surface levels of CSF1R (F_(1,28)_=7.72, *p*=0.01), CD11b (F_(1,28)_=137.6, *p*<0.0001), and CD68 (F_(1,28)_=102.2, *p*<0.0001), as well as decreased CX3CR1 expression (F_(1,28)_=244.4, *p*<0.0001). Moreover, microglia from *P2ry12-/-* mice showed higher side scatter (SSC), suggesting an increase in intracellular complexity (F_(1,28)_=7.91, *p*=0.009). In addition, CUS increased microglial CSF1R (*p*=0.0001) and SSC (*p*=0.006) in *P2ry12-/-* mice. These findings indicate that loss of P2Y12 leads to significant alterations in microglial phenotype and responsivity to CUS.

### Loss of P2Y12 causes dysregulation of microglial phagocytosis in the medial prefrontal cortex and prevents microglia-mediated dendritic remodeling in chronic stress

To examine neuron-microglia interactions, *P2ry12-/-* and wild-type mice were crossed with Thy1-GFP(M) mice. Following 14 days of CUS brains were processed for confocal microscopy (Fig.3A, Fig.S5-6). Similar to prior work, exposure to CUS increased the number of FosB+ cells/mm^2^ in the mPFC regardless of genotype (F_(1,26)_= 46.54, *p*<0.0001; Fig.3B, Fig.S5). Interestingly, *P2ry12-/-* mice had increased microglial density (IBA1+ cells/mm^2^; F_(1,26)_=83.95, *p*<0.0001, Fig.3C) and clustering (Fig.S5) in the mPFC. Loss of P2Y12 had little effect on basal levels of IBA1+ area per microglia in the mPFC (Fig.3D). However, CUS increased microglial area in the mPFC in wild-type, but not *P2ry12-/-* mice (F_(1,26)_=11.27, *p*=0.002).

**Figure 3.**
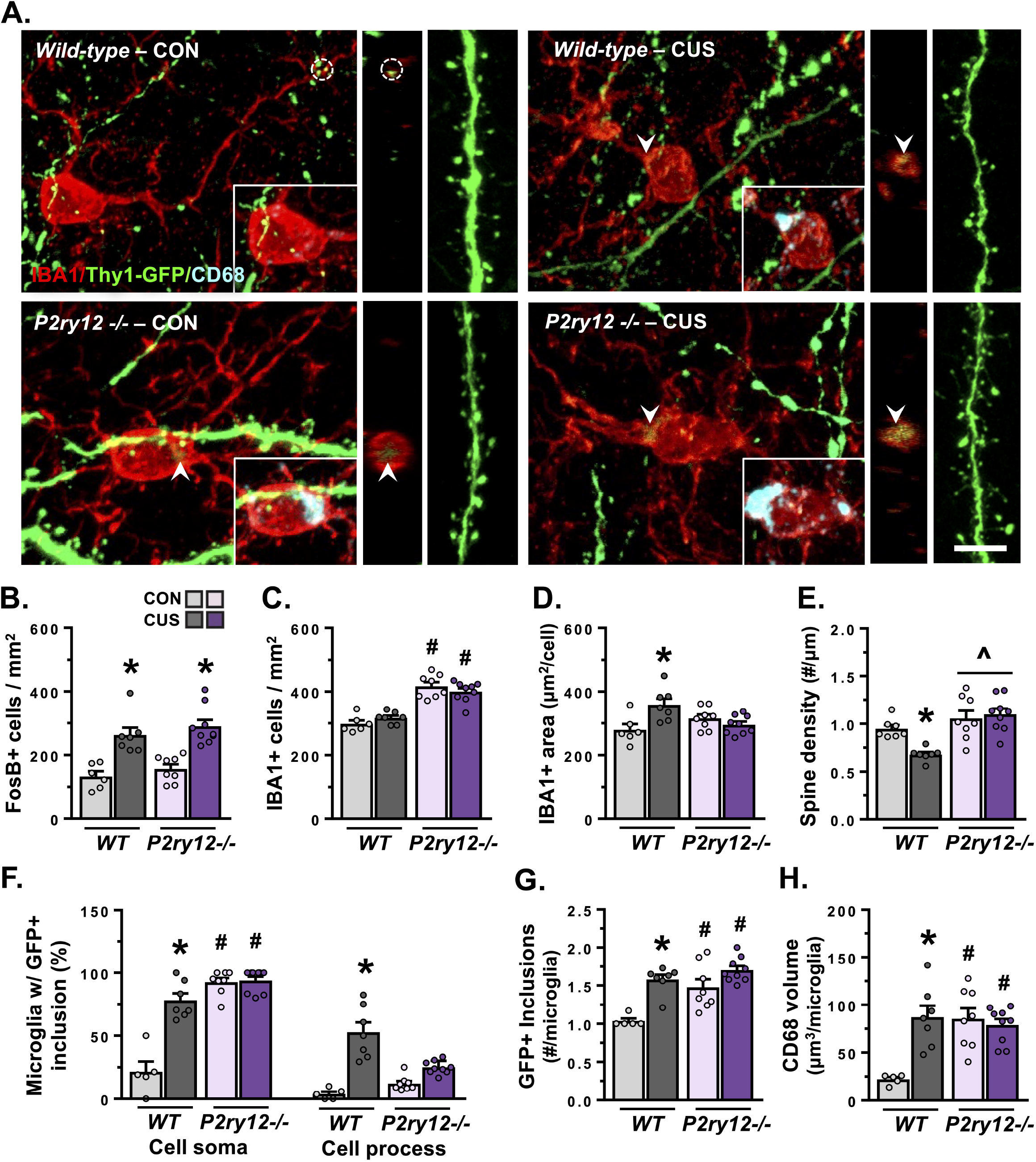
Loss of P2Y12 increases microglial phagocytosis in the medial prefrontal cortex and prevents microglia-mediated dendritic remodeling in chronic stress. Male Thy1-GFP(M) wild-type or *P2ry12*-/- mice were exposed to 14 days of chronic unpredictable stress (CUS) or were handled as controls. Approximately 4 hours after the final stressor, mice were perfused and brains were collected, sectioned, immunostained, and imaged (*n =* 5-9/group). **A**. Confocal images of microglia (IBA1, red), dendritic segments (Thy1-GFP, green), and microglial lysosomes (CD68, blue, inset) were obtained from lamina I of the mPFC. Alongside merged channels, an orthogonal cross-section (matching the noted location) and representative dendritic segment is depicted for each group. Microglial processes in close proximity to dendritic elements are noted within dashed circles, arrows indicate a dendritic element localized within a microglial cell body or process extension. White scale bar represents 5 μm. See Fig.S6 for additional images. **B**. Average number of FosB+ cells/mm^2^ in the mPFC. **C**. Average number of IBA1+ cells/mm^2^ in the mPFC. **D**. Average IBA1+ area per microglial cell. **E**. Average dendritic spine density. **F**. Proportion of microglia with GFP+ inclusions in the cell soma (left) and in cell processes (right). **G**. Number of GFP+ inclusions within microglia with dendritic elements. **H**. Average volume of CD68 per microglial cell. Bars represent mean ± S.E.M. ^*^ *p*<0.05 compared to same-genotype unstressed animal. ^#^ *p*<0.05 compared to unstressed wild-type animal. ^ p<0.05 main effect of genotype.

Assessment of dendritic compartments showed that exposure to CUS reduced spine density on apical dendritic arbors in layer I of the mPFC in wild-type mice (F_(1,27)_=7.00, *p*=0.01, Fig.3A&E). *P2ry12-/-* mice overall showed a modest increase in dendritic spine density (F_(1,27)_=20.20, *p*=0.0001) and this was unaffected by CUS. As expected, we found that CUS increased the proportion of microglia with GFP+ inclusions in wild-type mice. Surprisingly, we also observed high levels of GFP+ inclusions in PFC microglia in both unstressed and stressed *P2ry12-/-* mice. To expand on this finding, we quantified the presence of GFP+ inclusions in the soma and processes of microglia in the PFC (Fig.3F). These analyses showed that CUS increased GFP+ inclusions in both the soma (F_(1,25)_=34.36, *p*<0.0001) and processes (F_(1,25)_=18.3, *p*=0.0002) of microglia in wild-type mice, but not in *P2ry12-/-* mice. The basal accumulation of GFP+ inclusions in *P2ry12-/-* mice was primarily found in the soma (F_(1,25)_=84.92, *p*<0.0001, Fig.3F). Regardless of cellular localization, both stress and genotype had significant effects on the number of GFP+ inclusions per phagocytic microglia (F_(1,25)_=4.28, *p*=.049, Fig.3G), and the volume of CD68+ material per microglia (F_(1,25)_=13.96, *p*=.001, Fig.3H) in the mPFC.

Prior studies suggest that accumulation of lysosomes in the cell body may lead to autofluorescent inclusions (i.e., lipofuscin) [16]. To test this, PFC microglia with CD68+ lysosomes were visualized in wild-type and *P2ry12-/-* mice lacking the Thy1-GFP(M) transgene. In wild-type mice there were few autofluorescent CD68+ lysosomes, however, autofluorescent material was observed in somatic lysosomes in *P2ry12-/-* mice (Fig.S7). The mean intensity of these somatic accumulations was significantly lower than that of inclusions observed in mice carrying the Thy1-GFP(M) transgene (Fig.S7). These results indicate that GFP+ inclusions in microglia are composed of neuronal material in wild-type Thy1-GFP(M) mice. In contrast, it appears that GFP+ inclusions in microglia from *P2ry12-/-* Thy1-GFP(M) animals are composed of both autofluorescent and neuronal material. Altogether, P2Y12 appears to play a critical role in establishing microglial density and phagocytosis in the mPFC regardless of stress. These data further suggest that P2Y12 is important in guiding stress effects on microglial morphology and microglia-neuron interactions in this region.

### Pharmacological blockade of microglial P2Y12 attenuates stress effects on behavior and shifts microglial phenotype in frontal cortex

To complement previous experiments, P2Y12 was manipulated pharmacologically using either clopidogrel, a P2Y12 antagonist that can access the brain, or ticagrelor, an antagonist which cannot cross the blood-brain barrier. Unstressed and CUS-exposed mice treated with either vehicle, clopidogrel, or ticagrelor were assessed in the FST and TOR (Fig.4A). CUS increased immobility in the FST (F_(2,72)_=3.20, *p*=0.046) in vehicle- (*p*=0.007) and ticagrelor-treated mice (*p*=0.02; Fig.4B). Drug treatment affected the time spent exploring objects in the TOR (F_(2,72)_=4.21, *p*=0.02), however, post-hoc comparisons uncovered no group level differences (Fig.4C). Similar to the FST, CUS caused deficits in temporal object recognition (F_(1,72)_=23.92, *p*<0.0001) which were only detected in vehicle-(*p*<0.0001) and ticagrelor-treated mice (*p*=0.005; Fig.4D). Exposure to CUS reduced weight gain in both vehicle- and clopidogrel-treated animals (Table.S3). Collectively, these findings indicate that blocking P2Y12 on microglia (i.e. clopidogrel) but not peripheral cells (i.e. ticagrelor) attenuates stress effects on behavior.

**Figure 4.**
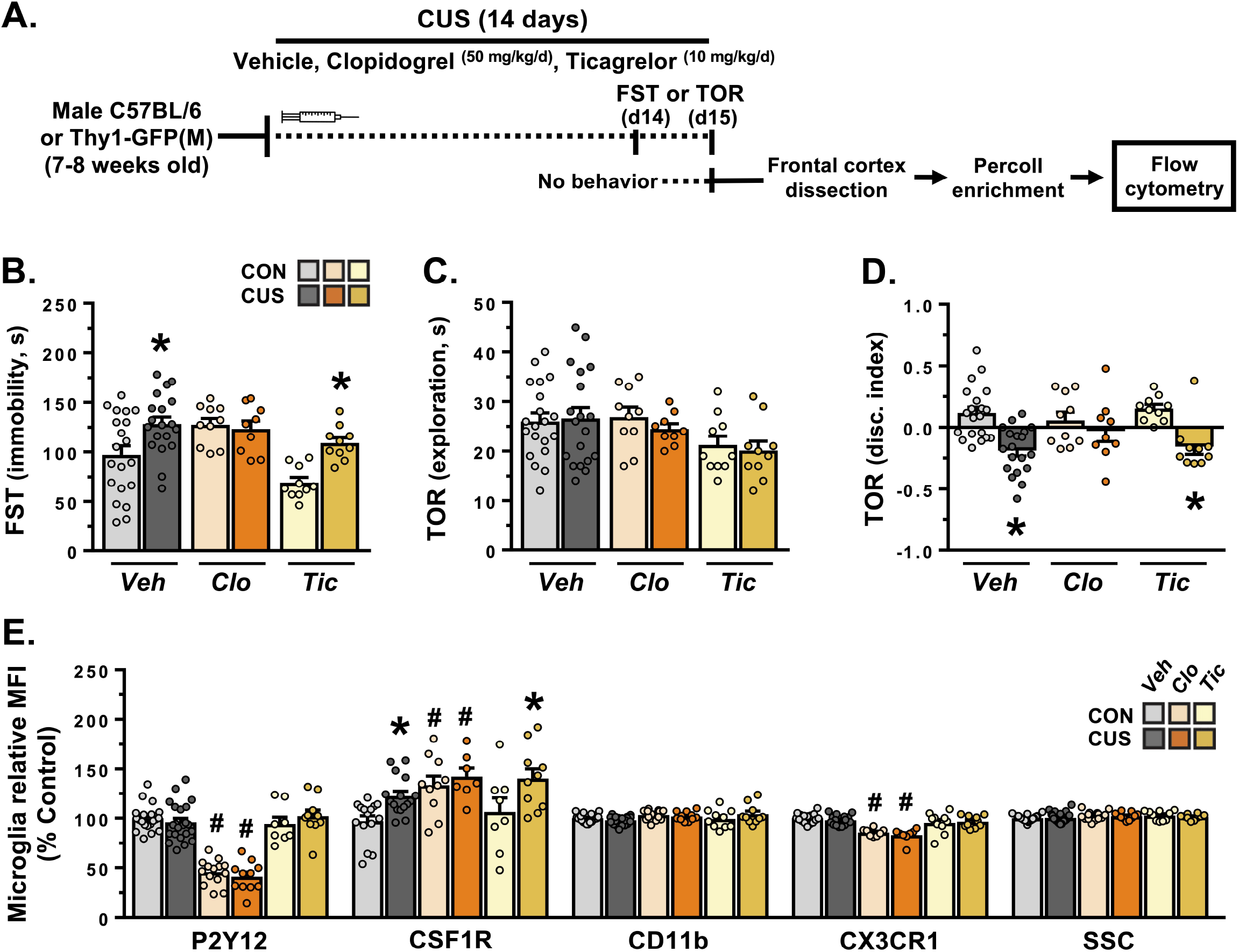
Pharmacological blockade of microglial P2Y12 attenuates stress effects on behavior and shifts microglial phenotype in the frontal cortex. **A**. Male wild-type or Thy1-GFP(M) mice were exposed to 14 days of chronic unpredictable stress (CUS) or were handled as controls. During this time, animals received daily injections of either vehicle, clopidogrel, or ticagrelor. One cohort was subjected to behavioral testing (*n =* 9-20/group). **B**. Average time spent immobile in the forced swim test (FST). **C**. Average time spent exploring objects in the temporal object recognition task (TOR). **D**. Discrimination index in the TOR. **E**. In a separate cohort of mice, brains were extracted, frontal cortex was dissected out, and microglia were isolated and characterized using flow cytometry (*n =* 7-22/group). Normalized mean fluorescence intensity of P2Y12, CSF1R, CD11b, and CX3CR1 in frontal cortex microglia. Normalized mean side scatter profile is shown. Bars represent mean ± S.E.M. ^*^ *p*<0.05 compared to same-treatment unstressed animal. ^#^ *p*<0.05 compared to unstressed vehicle-treated animal.

In a separate cohort, we analyzed the phenotype of frontal cortex microglia using flow cytometry (Fig.4E). Similar to genetic loss, treatment with clopidogrel reduced expression of P2Y12 (F_(2,79)_=98.59, *p*<0.0001) and CX3CR1 (F_(2,62)_=30.39, *p*<0.0001). There was an overall effect of treatment on microglial CD11b (F_(2,83)_=3.39, *p*=0.04); however, no group level differences emerged with post-hoc comparisons. Increased levels of microglial CSF1R were observed in mice treated with clopidogrel (F_(2,60)_=5.96, *p*=0.004) and those exposed to CUS (F_(1,60)_=10.66, *p*=0.002). Group comparisons showed that CUS increased microglial CSF1R in both vehicle- (*p*=0.03) and ticagrelor-treated mice (*p*=0.03), but not clopidogrel-treated mice. Neither drug treatment nor CUS had an effect on microglial SSC. Thus, pharmacological blockade of P2Y12 caused changes in microglial phenotype that resembled genetic loss, while also preventing stress effects on CSF1R expression in frontal cortex microglia.

### Inhibiting microglial P2Y12 blocks stress-induced phagocytosis of dendritic elements and subsequent dendritic spine loss in the mPFC

We next analyzed neuronal function, microglial phenotype, and microglia-neuron interaction in the mPFC using confocal microscopy (Fig.5A-J). CUS increased the number of FosB+ cells/mm^2^ in the mPFC in both vehicle- and clopidogrel-treated animals (F_(1,26)_=24.18, *p*<0.0001; Fig.5B). Neither CUS nor treatment with clopidogrel affected the number of IBA1+ cells/mm^2^, P2Y12+ cells/mm^2^, or microglial clustering in the mPFC (Fig.5C, Fig.S8). However, CUS increased microglial morphological area, and this was blocked by treatment with clopidogrel (F_(1,26)_=13.59, *p*=0.001; Fig.5D). Similar to our flow cytometry findings, treatment with clopidogrel reduced microglial P2Y12 intensity in the PFC (F_(1,26)_=22.59, *p*<0.0001; Fig.5E).

**Figure 5.**
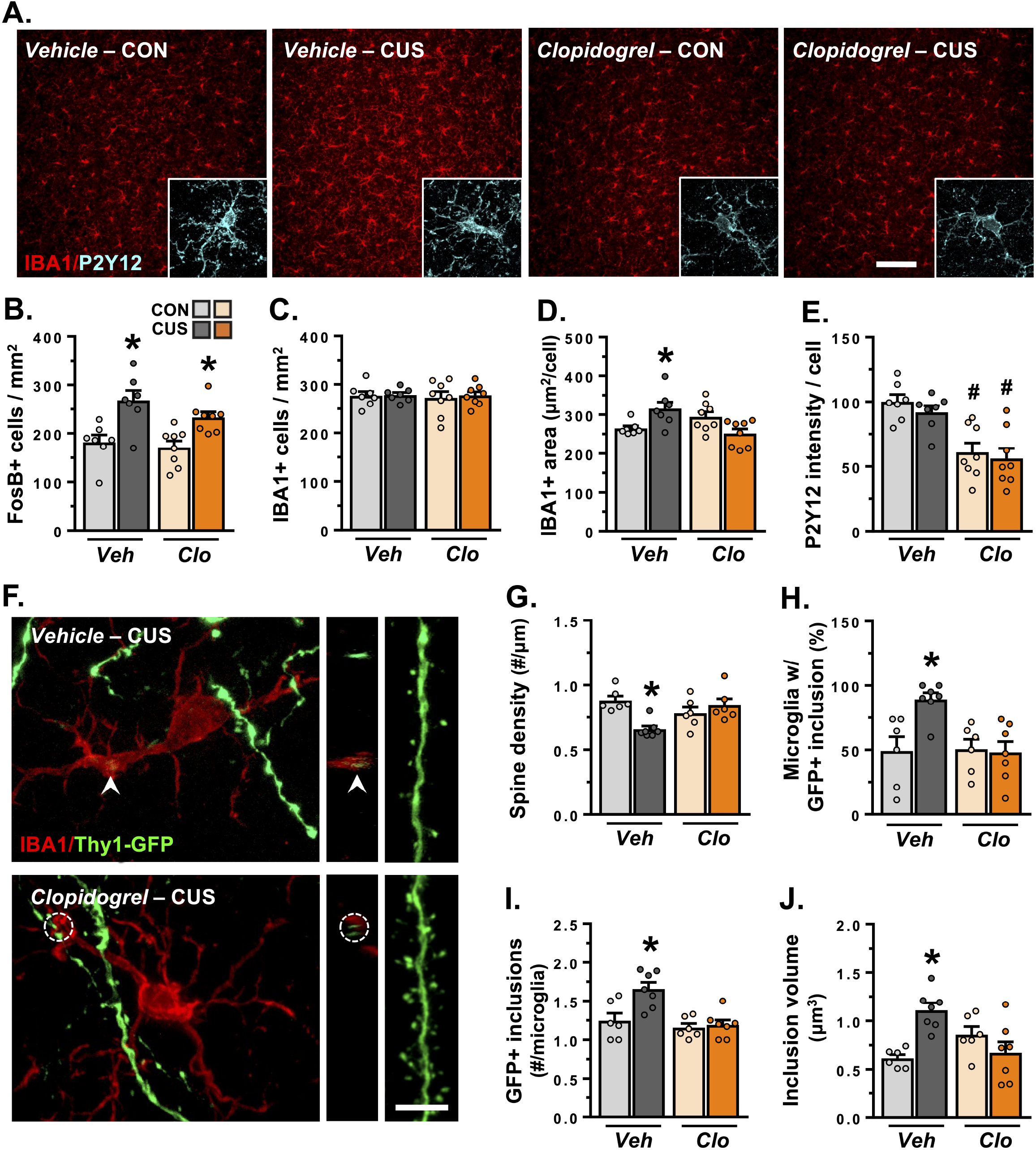
Inhibiting microglial P2Y12 blocks stress-induced phagocytosis of dendritic elements and subsequent dendritic spine loss in the mPFC. Male Thy1-GFP(M) mice were exposed to 14 days of chronic unpredictable stress (CUS) or were handled as controls. During this time, animals received daily injections of either vehicle or clopidogrel. Approximately 4 hours after the final stressor, mice were perfused and brains were collected, sectioned, immunostained, and imaged (*n* = 6-8/group). **A**. Confocal images of microglial IBA1 (red) and P2Y12 (cyan) in the mPFC. White scale bar represents 100 μm. **B**. Average number of FosB+ cells/mm^2^ in the mPFC. **C**. Average number of IBA1+ cells/mm^2^ in the mPFC. **D**. Average IBA1+ area per microglial cell. **E**. Average intensity (A.U.) of P2Y12 staining per microglial cell (relative to vehicle treated control group). **F**. Confocal images of microglia (IBA1, red) and dendritic segments (Thy1-GFP, green) were obtained from lamina I of the mPFC. Alongside merged channels, an orthogonal cross-section (matching the noted location) and representative dendritic segment is depicted for groups exposed to CUS. Microglial processes in close proximity to dendritic elements are noted within dashed circles, arrows indicate a dendritic element localized within a microglial cell body or process extension. White scale bar represents 5 μm. See Fig.S9 for additional images. **G**. Average dendritic spine density. **H**. Proportion of microglia with GFP+ inclusions. **I**. Number of GFP+ inclusions within microglia with dendritic elements. **J**. Average GFP+ inclusion volume per phagocytic microglia. ^*^ *p*<0.05 compared to same-treatment unstressed animal. ^#^ *p*<0.05 compared to unstressed vehicle-treated animal.

Blocking microglial P2Y12 with clopidogrel prevented spine loss on apical dendrites in the mPFC of CUS-exposed mice (F_(1,21)_=12.98, *p*=0.002; Fig.5F-G, Fig.S8). In line with this, CUS increased the proportion of microglia with GFP+ inclusions in the mPFC in vehicle-, but not clopidogrel-treated animals (F_(1,22)_=6.76, *p*=0.02; Fig.5H). Further analyses indicate that treatment with clopidogrel prevented CUS-induced increases in the number of GFP+ inclusions (F_(1,22)_=5.61, *p*=0.03; Fig.5I), the average volume of GFP+ inclusions (F_(1,22)_=15.94, *p*=0.001; Fig.5J), and the maximum GFP+ inclusion volume per phagocytic microglia in the PFC (Fig.S8-9).

## Discussion

Various stress-linked psychiatric disorders, including MDD, are marked by dendritic atrophy and synapse loss in the PFC. Recent preclinical studies suggest that microglia contribute to stress-induced changes in dendritic structure and associated behavioral consequences, however, the pathways that direct microglia-neuron interactions in this context remain unclear. Our primary findings indicate that P2Y12 signaling is required for stress-induced microglial phagocytosis of neuronal elements and mediates spine loss on pyramidal neurons in the PFC, which underlies deficits in working memory and stress-coping behavior. Our data also demonstrate that P2Y12 is essential for establishing microglial phenotype and prefrontal architecture, in the absence of stress.

### P2Y12 is a critical regulator of microglial phenotype and function

Studies have identified P2Y12 as a critical receptor in the microglial ‘sensome’, guiding microglial surveillance and physical microglia-synapse contact [6, 8, 17, 18]. In line with these findings, our data show that P2Y12 broadly regulates microglial phenotype and function in the frontal cortex. Genetic loss or pharmacological blockade of P2Y12 shifted the homeostatic phenotype of microglia with increased surface protein levels of CSF1R and CD11b, and decreased expression of CX3CR1. This is relevant because CSF1R and CD11b have been shown to drive microglia-neuron interaction and synaptic remodeling [3, 19], whereas CX3CR1-signaling can either encourage- or suppress-aspects of microglial function [20, 21]. These phenotypic alterations are likely a compensatory mechanism through which microglia attempt to survey and modulate synaptic structures with deficient P2Y12 signaling.

Another important finding is that genetic loss, but not pharmacological blockade, of P2Y12 increased the density and morphology of microglia in the PFC of unstressed adult mice. In addition, microglia in unstressed *P2ry12-/-* mice accumulated large somatic CD68+ inclusions that included autofluorescent and neuronal material. This suggests that these microglia have impaired regulation of lysosomes and possibly increased levels of phagocytosis or autophagy. Future studies will be needed to determine the material in these inclusions and mechanisms that lead to their accumulation. Developmental loss of P2Y12 also increased the number of dendritic spines on apical arbors in this region, suggesting alterations in synaptic plasticity. These neurobiological effects may be due to impaired microglia function as P2Y12 guides rearrangement of the microglial landscape [17]. Other studies have established that microglial P2Y12 mediates experience-dependent synaptic plasticity in the developing visual cortex [8]. Thus, genetic loss of P2Y12 may cause a discordant microglia phenotype that compromises microglia translocation and microglia microglia-mediated synaptic sculpting in the PFC, leading to microglial clustering and an overabundance of synapses in this region [22]. Despite these robust neurobiological effects, no behavioral alterations were detected in mice lacking P2Y12. This is consistent with a recent report showing only modest shifts in cognition and behavior in *P2ry12-/-* mice [23].

### Protracted neuronal activation shifts microglia to a ‘stress-like’ phenotype and disrupts working memory

Psychological stress increases neuronal activity across various corticolimbic brain regions, including the PFC. Microglia are uniquely attuned to neuronal activity, with various activation-associated signals (e.g. ADP/ATP) leading to functional adaptations in microglia. We’ve shown that blocking chronic stress-induced increases in neuronal activity using diazepam prevents microglia-mediated dendritic remodeling in the PFC and subsequent behavioral consequences [24]. One limitation of that study was that the pharmacological approach affected neurons across stress neurocircuitry. Here, we used direct chemogenetic manipulation to examine if repeated activation of pyramidal neurons in the mPFC promotes phenotypic changes in microglia that resemble those caused by CUS. CNO-treated mice showed robust expression of the activation marker FosB, indicating that this DREADD approach activated neurons in the PFC. These mice exhibited little change in stress-coping behavior as assessed in the FST. In contrast, mice treated with CNO had impaired temporal object recognition. This suggests that these tests engage different PFC pathways. Indeed, other studies have linked behavioral responses to FST with PFC projections to the dorsal raphe nucleus [25], whereas temporal object recognition appears to depend heavily on PFC connections with the mediodorsal thalamus and hippocampus [26, 27]. In line with this, FosB levels in the PFC were highly correlated with TOR performance, suggesting an activity-dependent link between prefrontal function and this working memory task. Protracted neuronal activity in the mPFC also caused changes in microglia density, morphology, and phenotype that resembled those caused by chronic stress [3, 11]. In particular, CNO-treated mice showed increased expression of several neuroimmune factors, including CD68, *Csf1r*, and *Cd11b* in the frontal cortex. These findings support the idea that neuronal activation in the mPFC provokes functional changes in microglia, including engagement of pathways important in microglia-synapse interaction.

### P2Y12 facilitates microglia-mediated dendritic remodeling, synapse loss, and behavioral dysfunction in stress

Stress-induced increases in neuronal activity alter glutamatergic neurotransmission and purine metabolism in the PFC [28, 29]. Prior work shows that glutamatergic signaling ‘attracts’ microglial processes via extracellular purines acting on microglial P2Y12 [7]. Several studies have shown that this signaling pathway promotes microglia-neuron interactions, which shape neuroplasticity and circuit function in various contexts [8, 10, 18, 30, 31]. In this study, CUS increased expression of the activation marker FosB in the PFC and disrupted both stress-coping behavior and working memory. Administration of the brain permeant P2Y12 antagonist clopidogrel attenuated these CUS-induced deficits, whereas treatment with ticagrelor, which cannot cross the blood-brain barrier, failed to rescue behavioral performance. Thus, blocking P2Y12 on microglia, specifically, reduces stress effects on behavioral and cognitive outcomes mediated by the PFC.

Consistent with prior studies, we found that CUS increased microglia-mediated remodeling of apical dendrites in the mPFC [2, 3]. Genetic loss or pharmacological blockade of microglial P2Y12 prevented microglia-mediated neuronal remodeling and attenuated synapse loss in the mPFC. Considering this with our data connecting neuronal activity and microglial function, it is plausible that CUS-induced neuronal activation of excitatory neurons in the PFC engages P2Y12 signaling in microglia, which increases the frequency of microglia-synapse interactions and subsequent microglia-mediated dendritic remodeling. This would be in line with other studies showing that P2Y12 is necessary for microglia to respond to fear- and seizure-associated activity in the hippocampus [18, 32], and that microglial P2Y12 can regulate experience-dependent synaptic sculpting [8]. It is important to note that the stress response is dynamic and likely engages other microglial pathways that are capable of modulating neuronal function [33]. Interestingly, a recent study showed that neuronal hyperactivity triggered a signaling cascade that involved P2Y12 and resulted in microglia providing negative regulatory feedback through release of the inhibitory molecule adenosine [30]. In this context, it is possible that P2Y12 signaling drives varied microglia functions across the duration of CUS. Additional studies will be needed to directly examine these temporal dynamics.

The preclinical findings presented here may be translationally relevant, as recent post-mortem analyses indicate heightened microglial *P2ry12, Cx3cr1*, and *Tmem119* expression in MDD [34, 35]. Intriguingly, these same studies found no differential expression of inflammation-associated molecules, suggesting that MDD is not marked by microglial ‘activation’ or neuroinflammation, but rather a more nuanced shift in microglial phenotype. This parainflammatory phenotype is observed in several models of chronic stress [36]. Given our data, it is interesting to speculate that microglial P2Y12 may play an important role in regulating synapse loss in MDD.

## Conclusion

Collectively, our findings demonstrate a role for neuronal activity and microglial P2Y12 in guiding stress effects on prefrontal structure, cognition, and behavior. These data may be clinically pertinent, as both prefrontal synapse loss and altered microglial *P2ry12* are implicated in MDD. This work is significant as it provides insight into pathways that direct microglia-neuron interactions, and how these interactions modulate synaptic structure and behavioral responses in stress-linked psychopathology.

## Supplementary Information

Supplementary information is available at BioRxiv’s website.

## Acknowledgments

The authors would like to thank Dr. Ania Majewska for kindly donating *P2ry12-/-* mice, and Dr. Grayson Sipe for advice regarding administration of clopidogrel and ticagrelor. The authors would also like to thank Dr. Lauren Vollmer for feedback on this manuscript. This work was supported by the National Institute of Mental Health (F32MH123051, JLB; R01MH123545, ESW) and the University of Cincinnati.

