## Supplemental Figures for "Microglial P2Y12 mediates chronic stress-induced synapse loss in the prefrontal cortex and associated behavioral consequences in male mice"

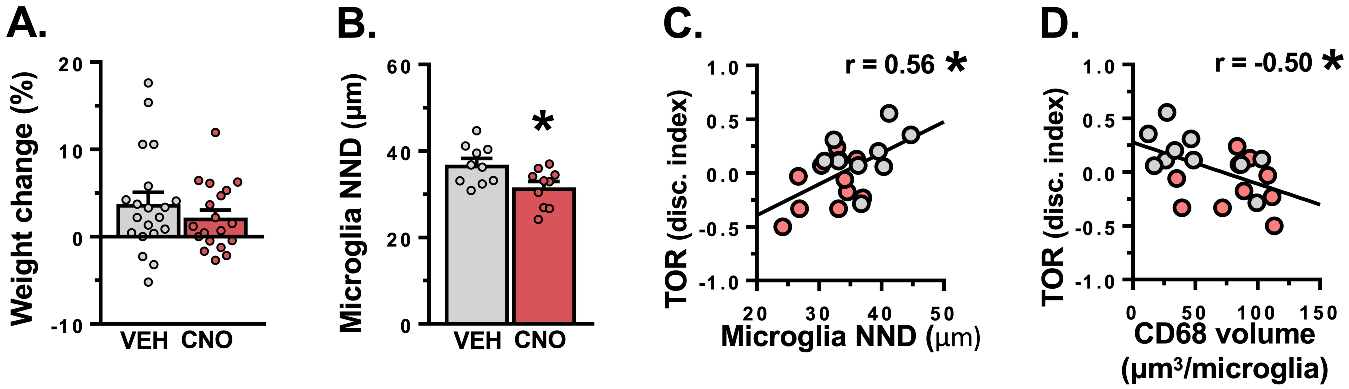


**Figure S1. Repeated neuronal activation induces microglial clustering in the medial prefrontal cortex and disrupts working memory performance.** Male hM3Dq-floxed mice received bilateral infusions of AAV5-CaMKIIa-mCherry-Cre into the mPFC. Post-recovery, animals received daily injections of either vehicle or CNO for 14 days (*n* = 9-10/group). **A.** Body weight was measured prior to CNO administration and post-CNO treatment, and percent weight change was calculated. Administration of CNO had no effect on animal weight gain. **B.** Microglial clustering was assessed using nearest neighbor distance (NND). Administration of CNO reduced the average microglial NND in the mPFC ($t_{(18)=}$2.66, *p*=0.02). **C-D.** Association between discrimination index in the TOR and (C) the average microglial NND (r=0.557, *p*=0.01) and (D) the average CD68+ lysosome volume per microglia in the mPFC (r=-0.503, *p*=0.02). Bars represent mean ± S.E.M. * *p*<0.05 compared to vehicle treated animal.

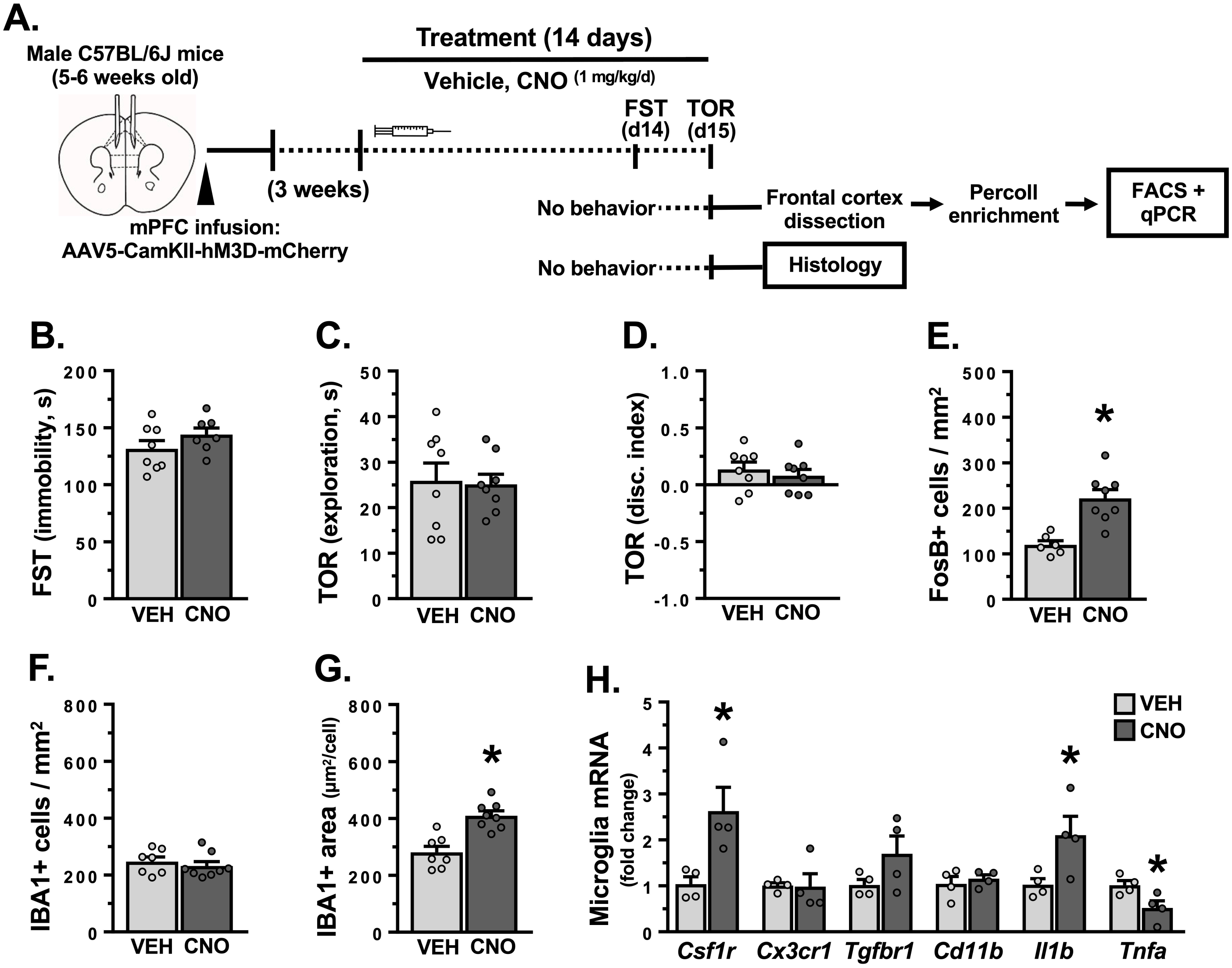


**Figure S2. Prolonged prefrontal activation driven by AAV5-CaMKIIα-hM3d-mCherry expression shifts microglial phenotype. A.** Male C57BL/6 mice received bilateral infusion of AAV5-CaMKIIα-hM3d-mCherry (DREADD) in the mPFC. Following recovery, mice were injected with vehicle or CNO (1 mg/kg/day, i.p.) for 14 days. One cohort was assessed in the forced swim test (FST) and temporal object recognition (TOR) test on subsequent days ($n$ = 8/group). Another cohort was used for immunohistology ($n$ = 8/group). Microglia were isolated and gene expression was analyzed in a separate cohort ($n$ = 4/group). **B.** Time spent immobile in the FST. **C.** Average time spent exploring objects in the TOR. **D.** Discrimination index in the TOR. **E.** Average number of FosB+ cells/mm$^2$ in the mPFC. **F.** Average number of IBA1+ cells/mm$^2$ in the mPFC. **G.** Average IBA1+ area per microglial cell. **H.** Following the final injection of CNO (24 h-post), frontal cortex was dissected and brain myeloid cells were enriched by Percoll gradient. Fluorescence activated cell sorting (FACS) was used to isolate microglia based on CD11b+/CD45lo expression. Following FACS, mRNA was collected from microglia. Relative fold change in gene expression in sorted frontal cortex microglia is shown. Bars represent mean ± S.E.M. * $p < 0.05$ compared to vehicle treated animals.

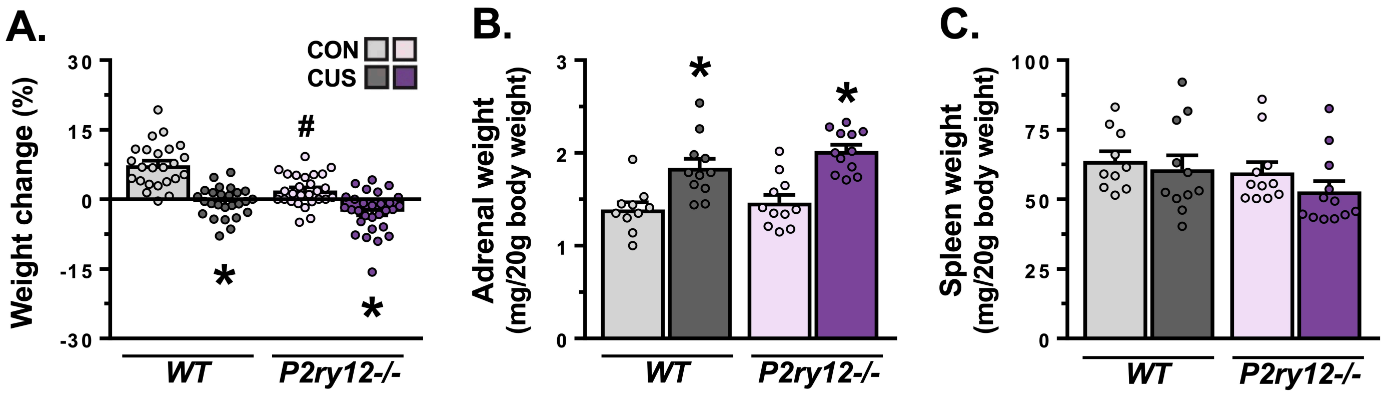


**Figure S3. Chronic unpredictable stress reduces weight gain and induces adrenal hypertrophy in male mice regardless of genotype.** Male wild-type or *P2ry12*-/- mice were exposed to 14 days of chronic unpredictable stress (CUS) or were handled as controls (CON). Body weight was measured prior to stress and post-CUS exposure, and percent weight change was calculated for all animals regardless of experimental endpoint (*n* = 24-29/group). In animals subjected to behavioral testing, stress-sensitive organ weight was measured (*n* = 10-12/group). **A.** Percent body weight change. CUS reduced weight gain regardless of genotype ($F_{(1,102)}$=4.80, $p$=0.03). **B.** Adrenal weight relative to animal body weight. CUS induced adrenal hypertrophy in both wild-type and P2ry12-/- animals ($F_{(1,40)}$=37.57, $p$<0.0001). **C.** Spleen weight relative to animal body weight. Bars represent mean ± S.E.M. * $p$<0.05 compared to same-genotype unstressed animal. # $p$<0.05 compared to unstressed wild-type animal.

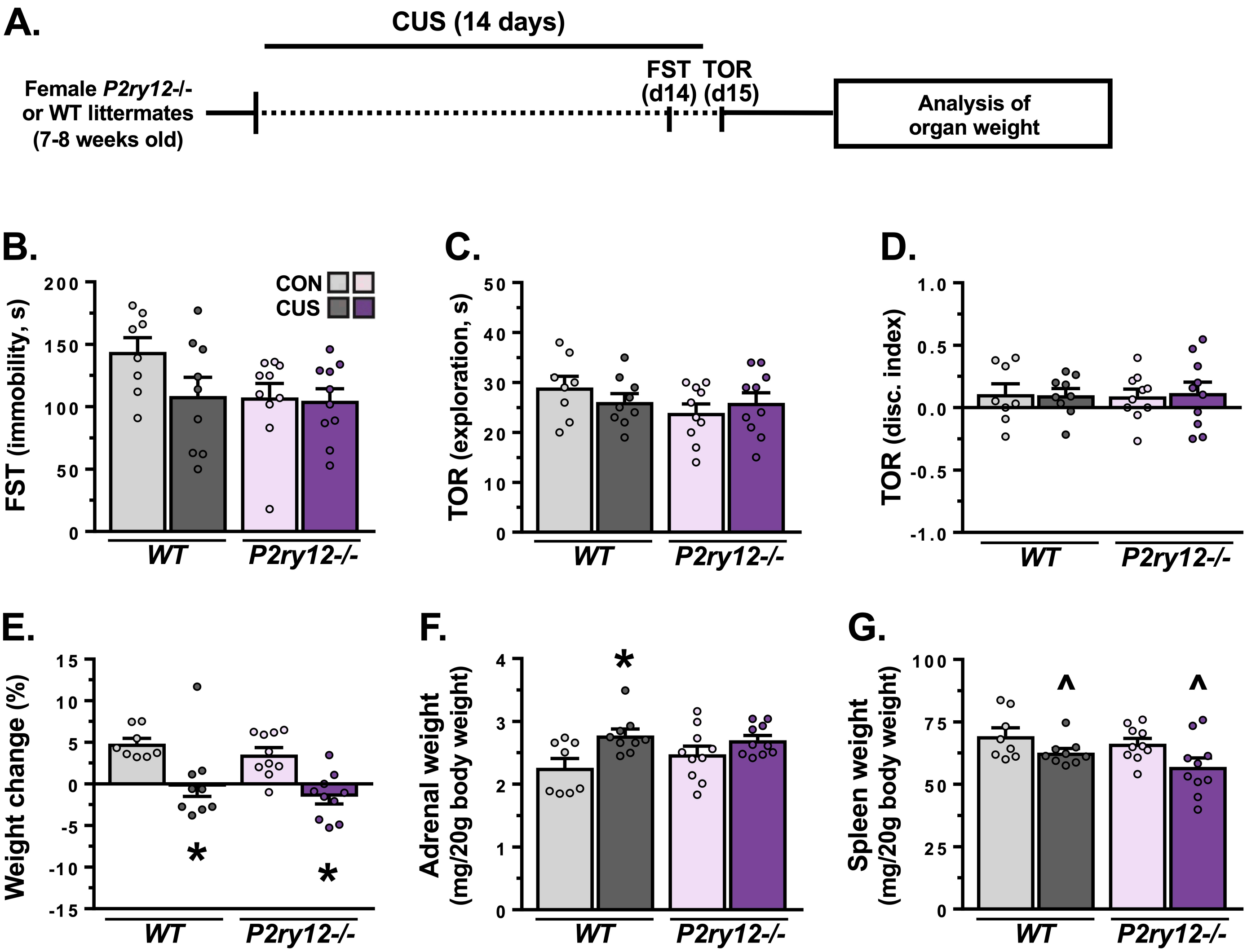


**Figure S4. Neither ablation of P2Y12 nor chronic unpredictable stress affected behavior in female mice, despite stress effects on physiology. A.** Female wild-type or *P2ry12-/-* mice were exposed to 14 days of chronic unpredictable stress (CUS) or were handled as controls (CON). Animals were subjected to behavioral testing followed by analysis of stress-sensitive organ weight ($n$ = 8-10/group). **B.** Average time spent immobile in the forced swim test (FST). **C.** Average time spent exploring objects in the temporal object recognition task (TOR). **D.** Discrimination index in the TOR. No effects of genotype or stress were detected for any of the behavioral endpoints measured ($p$>0.05). **E.** Body weight was measured prior to stress and post-CUS exposure, and percent weight change was calculated. CUS reduced weight gain ($F_{(1,33)}$=22.10, $p$<0.001) in both wild-type ($p$=0.01) and *P2ry12-/-* mice ($p$=0.003). **F-G.** Adrenal and spleen were dissected out, weighed, and normalized to animal body weight. CUS induced adrenal hypertrophy ($F_{(1,33)}$=10.26, $p$=0.003) and atrophy of the spleen ($F_{(1,33)}$=7.918, $p$=0.008). However, group-level differences in adrenal weight were only detected in wild-type animals ($p$=0.01), with no group-level differences detected in spleen weight. Bars represent mean ± S.E.M. $^*$ $p$<0.05 compared to same-genotype unstressed animal. $^\wedge$ $p$<0.05 main effect of CUS.

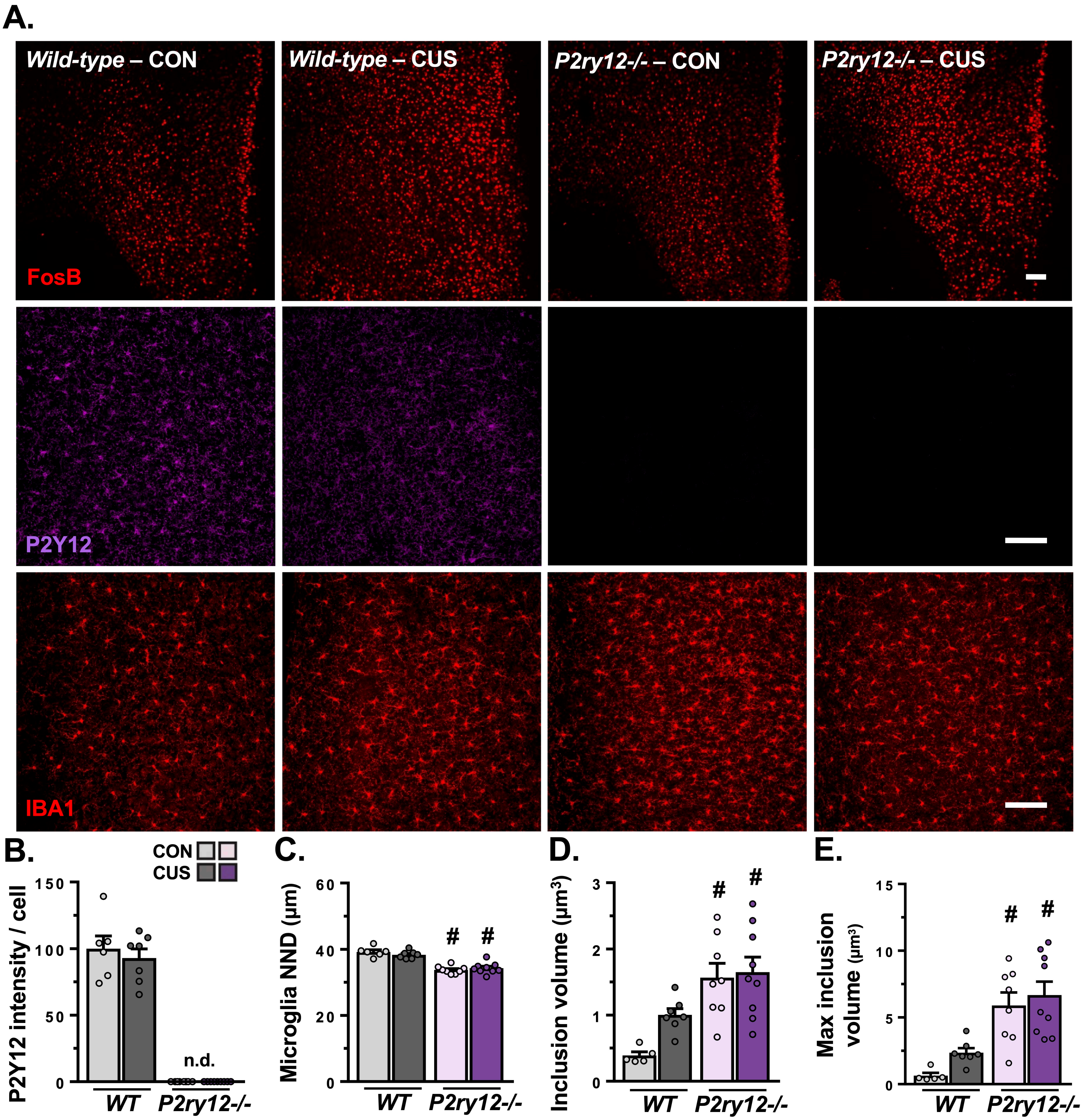


**Figure S5. Loss of P2Y12 increases microglial clustering in the medial prefrontal cortex.** Male wild-type or *P2ry12*-/- mice were exposed to 14 days of chronic unpredictable stress (CUS) or were handled as controls (CON). Approximately 4 hours after the final stressor, mice were perfused and brains were collected, sectioned, immunostained, and imaged ($n$ = 6-9/group). **A.** Confocal images depicting FosB, P2Y12, and IBA1 immunofluorescence in the mPFC are shown for each group. White scale bar represents 100 µm. **B.** P2Y12 intensity per microglial cell in the mPFC (relative to unstressed wild-type). P2Y12 was not detected (n.d.) in *P2ry12-/-* mice ($F_{(1,26)}$=381.9, $p$<0.0001). **C.** Microglial clustering was assessed using nearest neighbor distance (NND). The average microglial NND was reduced in P2ry12-/- mice ($F_{(1,26)}$= 85.55, $p$<0.0001). **D-E.** Graphs depict the average and maximum GFP+ inclusion volume detected in microglia in the mPFC. Bars represent mean ± S.E.M. # $p$<0.05 compared to unstressed wild-type animal.

6A.

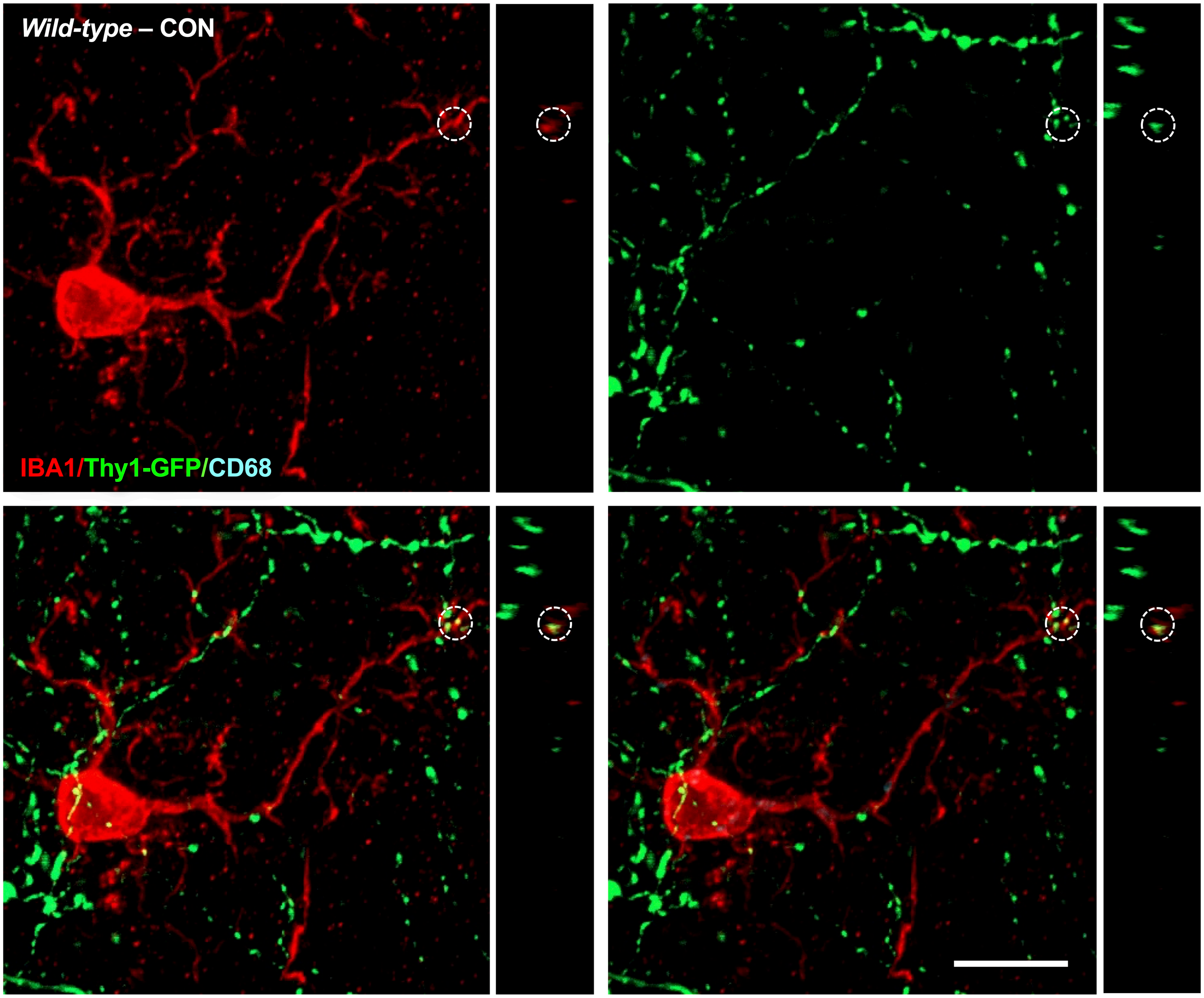


**Figure S6. Representative images of microglia-neuron interaction in the medial prefrontal cortex of wild-type and P2Y12 deficient mice.** Male Thy1-GFP(M) wild-type or *P2ry12-/-* mice were exposed to 14 days of chronic unpredictable stress (CUS) or were handled as controls. Approximately 4 hours after the final stressor, mice were perfused and brains were collected, sectioned, immunostained, and imaged. **A-D.** Confocal images of microglia (IBA1, red), dendritic segments (Thy1-GFP, green), and microglial lysosomes (CD68, blue) were obtained from lamina I of the mPFC. Each figure panel shows isolated channels for IBA1 (top left) and Thy1-GFP (top right), and merged channels for IBA1+Thy1-GFP (bottom left) and IBA1+Thy1-GFP+CD68 (bottom right). Alongside each individual and merged channel, an orthogonal cross-section (matching the noted location) is shown. Microglial processes in close proximity to dendritic elements are noted within dashed circles, arrows indicate a dendritic element localized within a microglial cell body or process. Not all microglial appositions or dendritic inclusions are marked. White scale bar represents 10 µm.

6B.

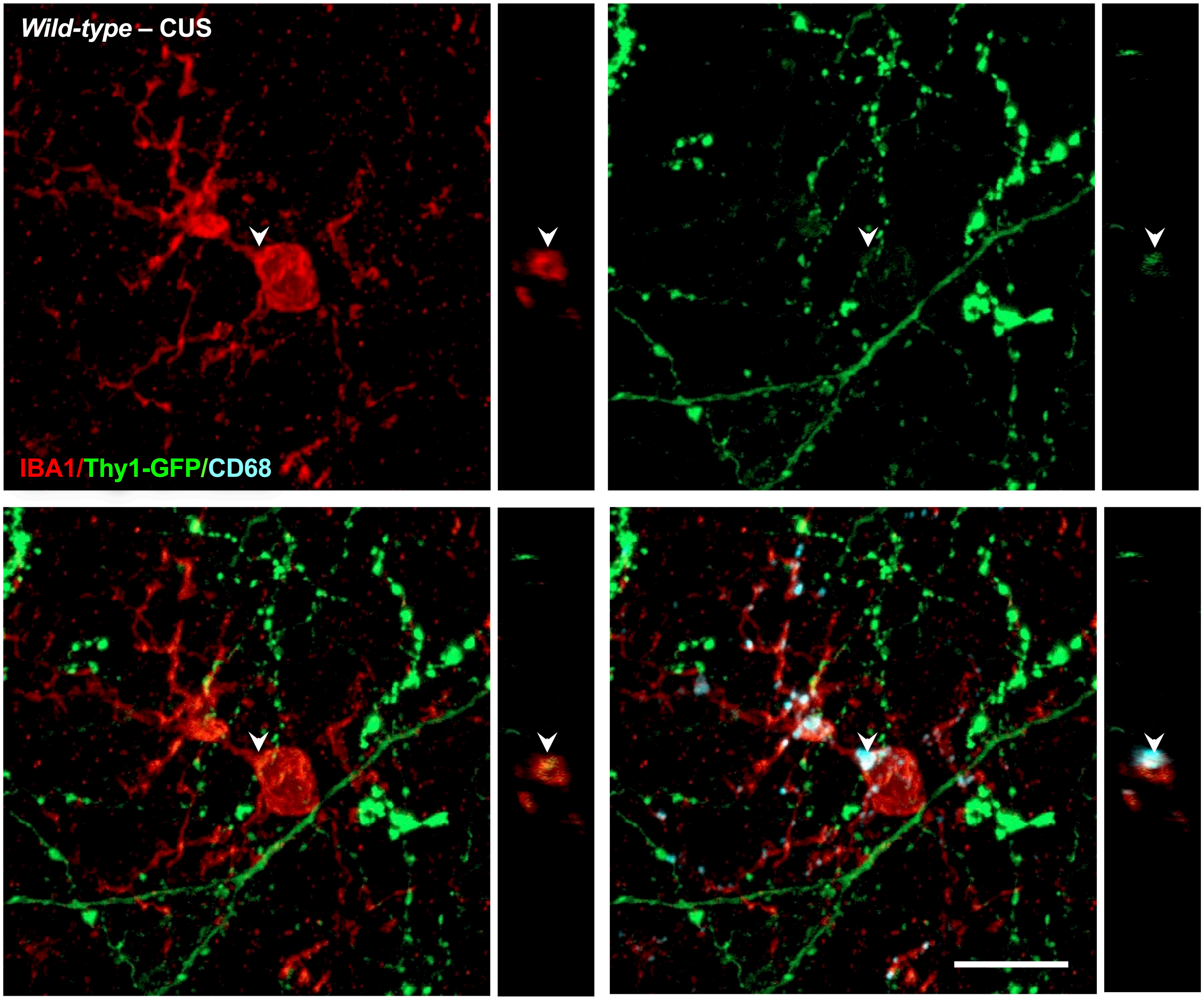

6C.

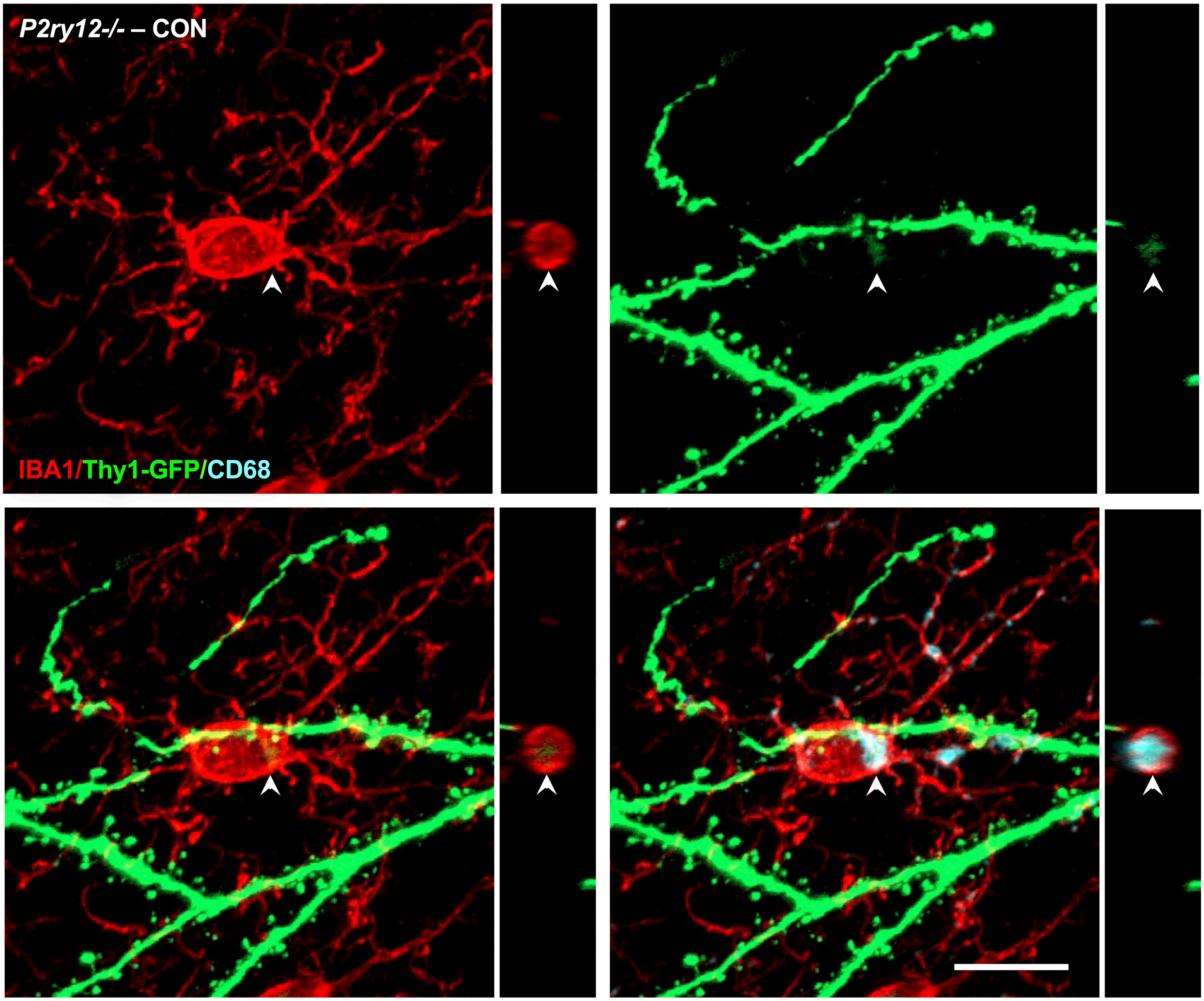

6D.

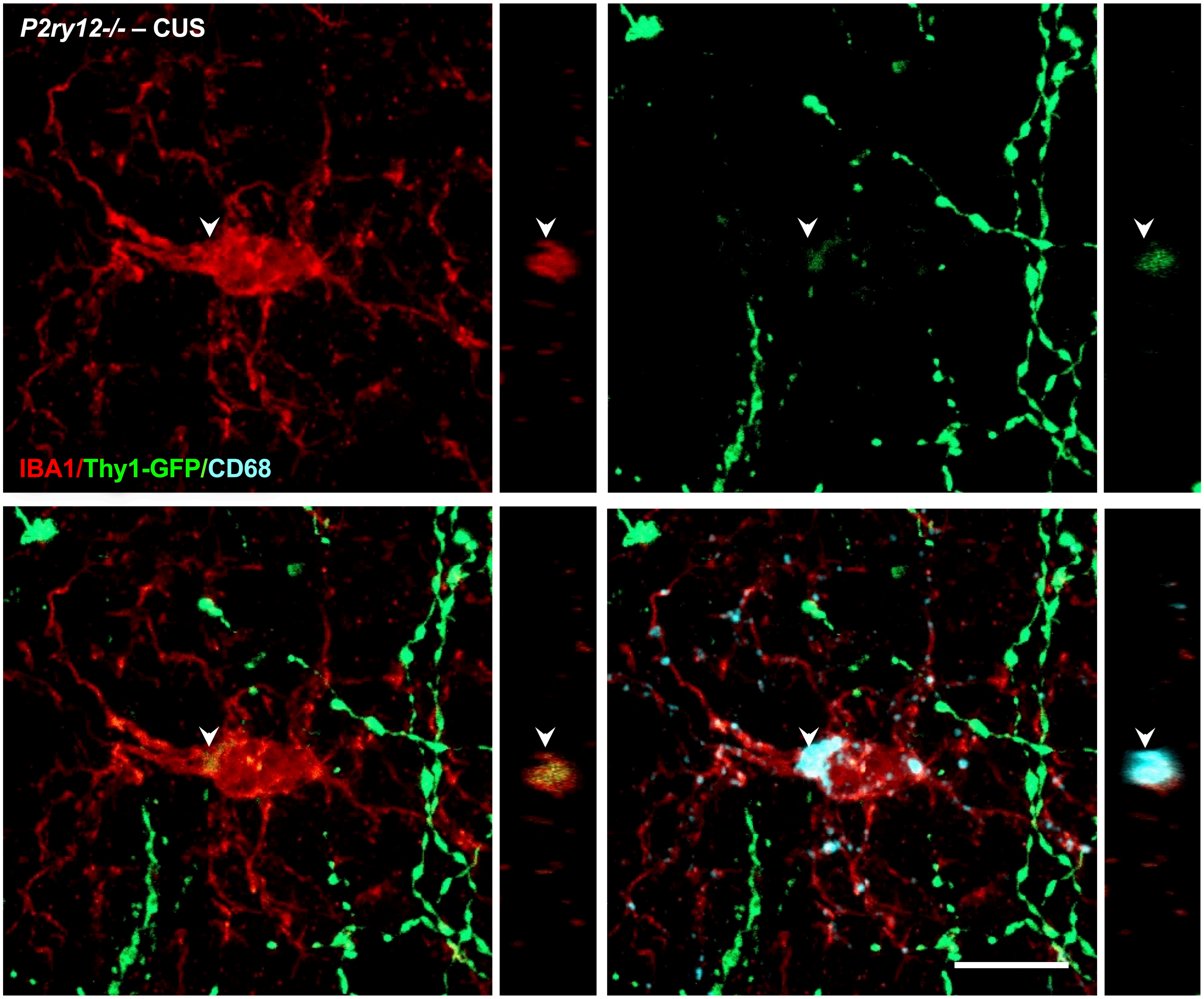

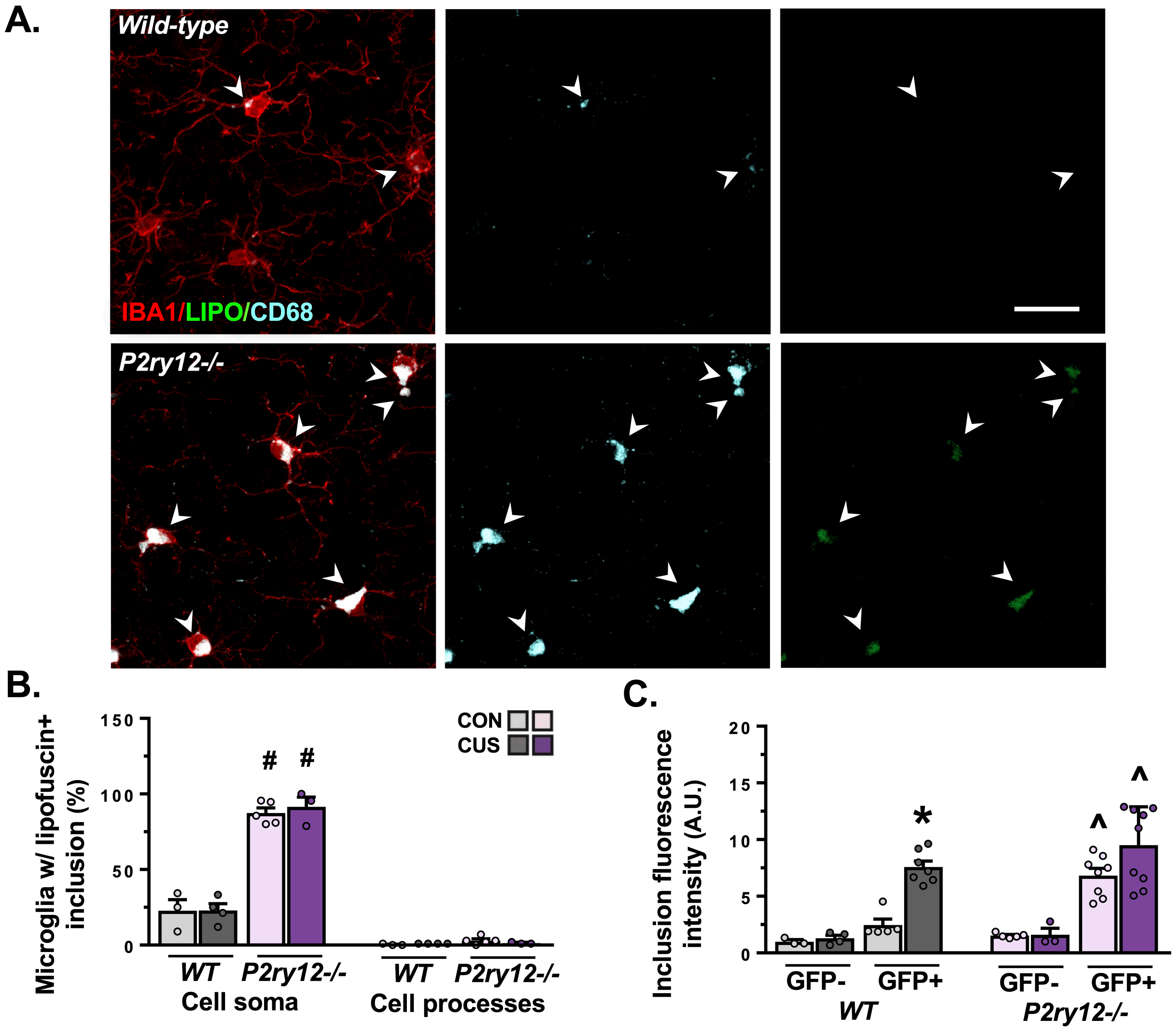


**Figure S7. Microglia in P2Y12 deficient mice accumulate autofluorescent lysosomal inclusions.** Male wild-type or *P2ry12-/-* mice lacking the Thy1-GFP(M) transgene were exposed to 14 days of chronic unpredictable stress (CUS) or were handled as controls. Brains were sectioned and stained for IBA1 and CD68. Autofluorescent lipid inclusions (e.g., lipofuscin) were imaged at 488 nm excitation with an emission spectra of 510–574 nm (*n* = 3-5/group). **A.** Confocal images of microglia (IBA1, red), microglial lysosomes (CD68, blue), and autofluorescent material (green) were obtained from lamina I of the mPFC. Image panels show merged channels for IBA1+CD68+lipofuscin (left) alongside isolated channels for CD68 (middle) and lipofuscin (right) in wild-type (top) and *P2ry12-/-* mice (bottom). Arrows indicate the location of various CD68+ lysosomes. White scale bar represents 20 µm. **B.** Proportion of microglia with lipofuscin+ inclusions in the cell soma (left) and in cell processes (right). A significantly greater proportion of microglia exhibited lipofuscin+ inclusions in the cell soma in *P2ry12-/-* mice ($F_{(1,11)}$=170.2, $p$<0.0001). Few lipofuscin+ inclusions were observed in microglial processes. **C.** Fluorescence intensity of inclusions was measured in both mice lacking the Thy1-GFP (M) transgene (*n* = 3-5/group) and Thy-GFP+ animals (*n* = 5-9/group; A.U. relative to wild-type animals lacking the Thy1-GFP(M) transgene). CUS increased the fluorescence intensity of inclusions exclusively in wild-type mice carrying the Thy1-GFP(M) transgene ($F_{(1,15)}$=20.49, $p$=0.0004). Inclusion intensity was significantly greater in *P2ry12-/-* mice carrying the Thy1-GFP (M) transgene as compared to GFP- animals ($F_{(1,21)}$=41.12, $p$<0.0001). Bars represent mean ± S.E.M. $^{\#}$ $p$<0.05 compared to unstressed wild-type animal. * $p$<0.05 compared to same-genotype unstressed animal. ^ p<0.05 compared to same-genotype animal lacking the Thy1-GFP(M) transgene.

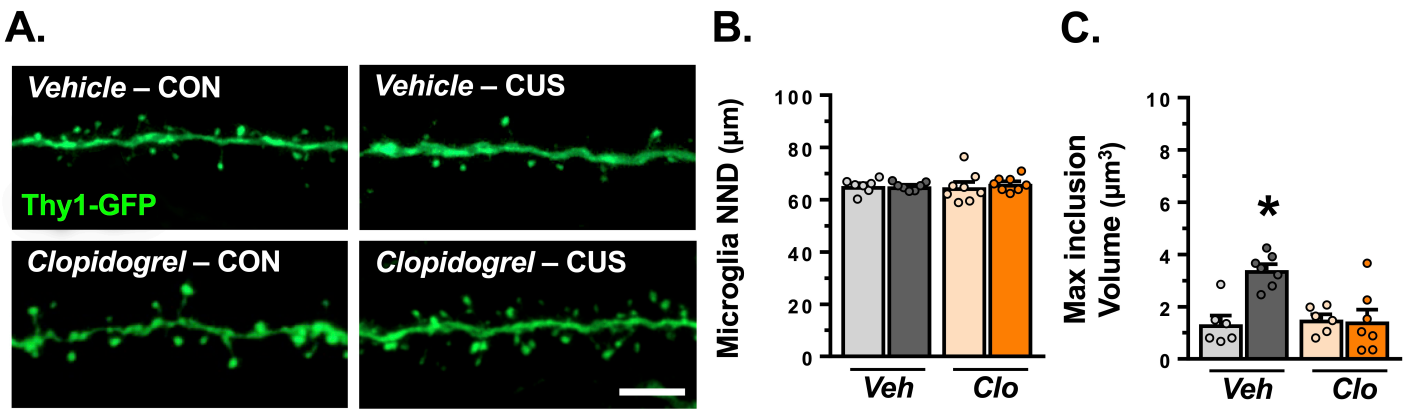


**Figure S8. Pharmacological blockade of microglial P2Y12 blocks stress-induced phagocytosis of dendritic elements and subsequent dendritic spine loss in the medial prefrontal cortex.** Male Thy1-GFP(M) mice were exposed to 14 days of chronic unpredictable stress (CUS) or were handled as controls. During this time, animals received daily injections of either vehicle or clopidogrel. Approximately 4 hours after the final stressor, mice were perfused and brains were collected, sectioned, immunostained, and imaged (n = 6-8/group). **A.** Representative images of dendritic segments (Thy1-GFP, green) from lamina I of the mPFC. White scale bar represents 5 µm. **B.** Microglial clustering was assessed using nearest neighbor distance (NND). Average microglial NND is shown. **C.** Graph depicts the maximum GFP+ inclusion volume detected in microglia in the mPFC. Administration of clopidogrel blocked CUS-induced increases in the maximum GFP+ inclusion volume detected in microglia ($F_{(1,22)}$=10.44, *p*=0.004). Bars represent mean ± S.E.M. * *p*<0.05 compared to same-treatment unstressed animal.

**Figure S9. (following pages). Representative images of microglia-neuron interaction in the medial prefrontal cortex of mice treated with either vehicle or clopidogrel.** Male Thy1-GFP(M) mice were exposed to 14 days of chronic unpredictable stress (CUS) or were handled as controls. During this time, animals received daily injections of either vehicle or clopidogrel. Approximately 4 hours after the final stressor, mice were perfused and brains were collected, sectioned, immunostained, and imaged. **A-D.** Confocal images of microglia (IBA1, red) and dendritic segments (Thy1-GFP, green) were obtained from lamina I of the mPFC. Alongside merged and individual channels, an orthogonal cross-section (matching the noted location) is shown for each experimental group. Microglial processes in close proximity to dendritic elements are noted within dashed circles, arrows indicate a dendritic element localized within a microglial cell body or process extension. White scale bar represents 10 µm.

9A.

9B.

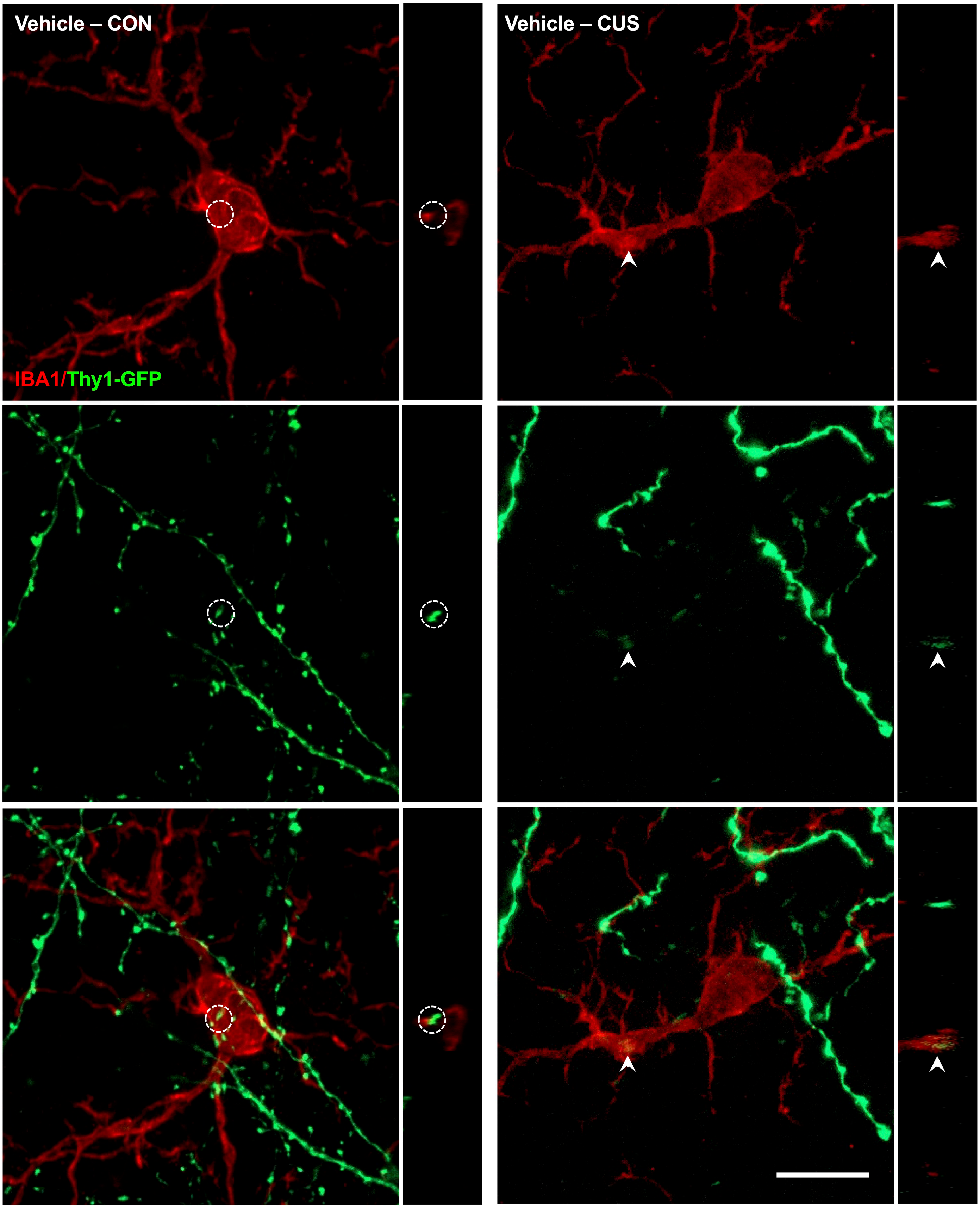

9C.

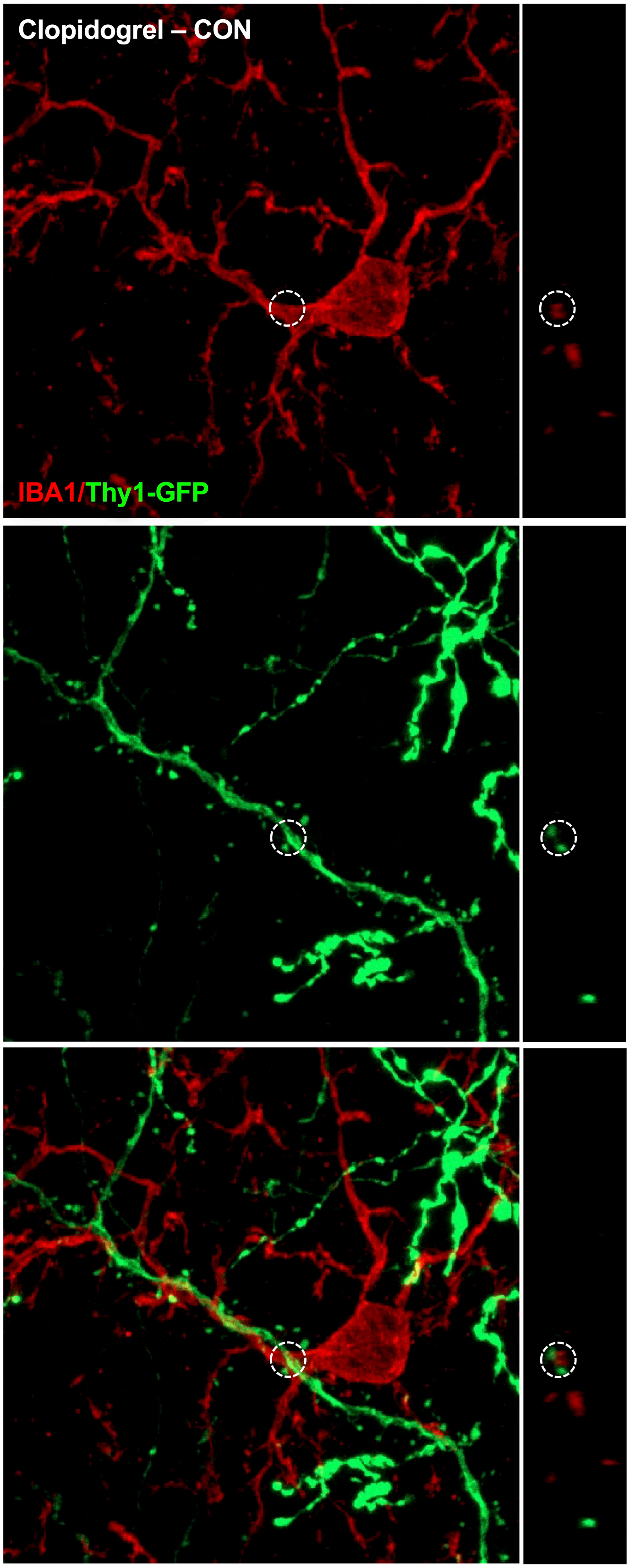


9D.

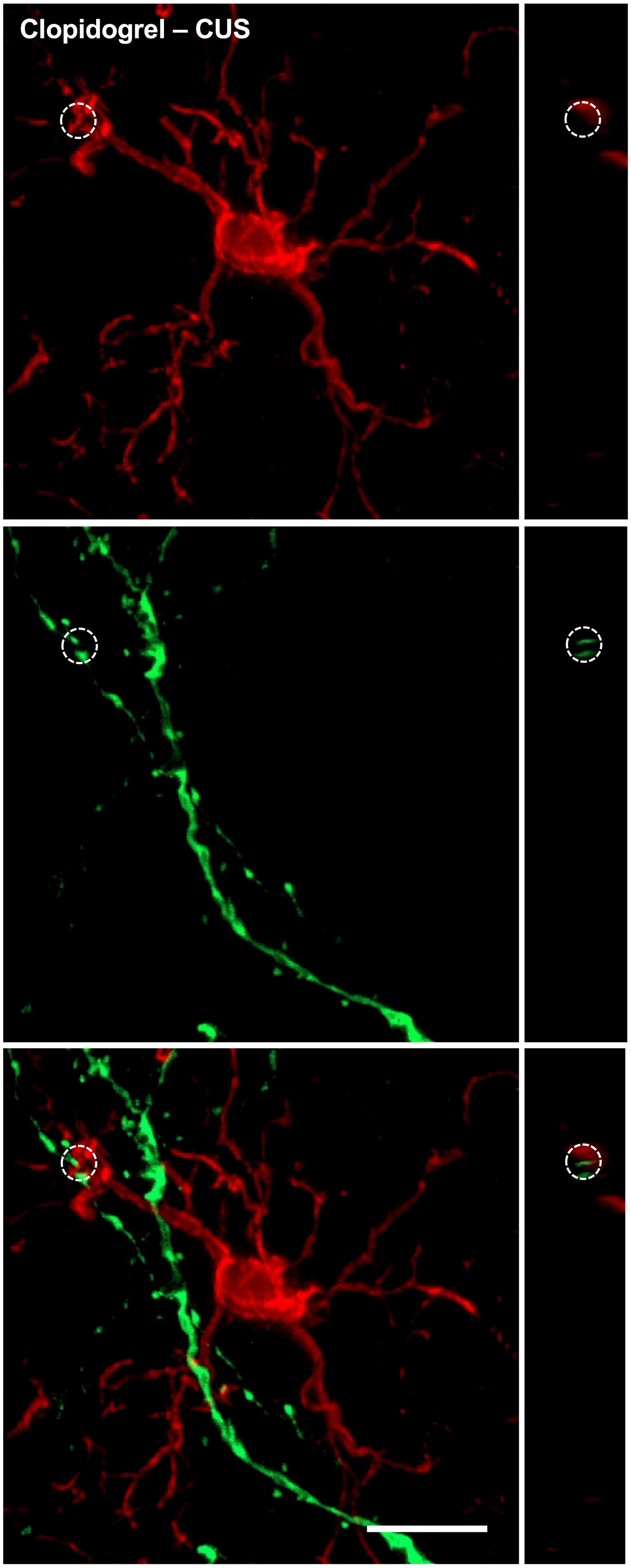
